# A biological-response compound representation allows chemical perturbation prediction across cell lines

**DOI:** 10.64898/2026.09.28.755146

**Authors:** Léa Kaufmann, Lola Le Breton, Elise Carraz-Billat, Quentin Fournier, Sébastien Lemieux

**Affiliations:** Institute for Research in Immunology and Cancer, Université de Montréal, Montréal, QC H3T 1J4, Canada; Department of biochemistry and molecular medicine, Université de Montréal, Montréal, QC H3T 1J4, Canada; Mila – Quebec AI Institute, Montréal, QC H2S 3H1, Canada; Polytechnique Montréal, Montréal, QC H3T 0A3, Canada; Université Paris-Saclay, Gustave Roussy, Inserm, UMR 1279 Tumor Cell Dynamics 94805 Villejuif, France; Department of computer science and operations research, Université de Montréal, Montréal, QC H3T 1N8, Canada

**Author notes:** Corresponding author: Sébastien Lemieux. These authors contributed equally to this work (Léa Kaufmann, Lola Le Breton).

## Abstract

Accurately predicting cellular responses to drugs remains a challenge with the potential to reduce experimental screening costs and accelerate drug discovery. Current computational approaches represent compounds through chemical structures, which carry little information regarding their activity within biological systems. We show that gene expression responses, measured in a reference cell line, can instead serve as transferable representations of chemical perturbations. The proposed framework, BioPert, uses biological representations with a small neural network to predict transcriptional delta responses in cell lines of different lineages. Across the Tahoe-100M and LINCS L1000 datasets, BioPert outperforms molecular fingerprints and embeddings learned from chemical structure, which provide limited improvements over random controls. BioPert’s prediction correlations reach 0.79 on Tahoe-100M, corresponding to an improvement of 0.34 over the next-best representation. On LINCS, performance varies with the experimental reproducibility of the test conditions, highlighting the impact of batch effects. Notably, BioPert considerably surpasses predictions obtained by copying the reference response, demonstrating that it captures context-dependent effects. Further analysis of a C32-cobimetinib case shows pathway-level accuracy of the predictions, including when the reference and target responses differ. These results open a new path for chemical perturbation prediction and could ultimately reduce the burden of phenotypic screening.

## 1 Introduction

Accurate computational models that predict cellular responses to chemical perturbations prior to *in vivo* experiments could play a fundamental role in accelerating phenotypic drug discovery and making drug repurposing more selective and scalable. This goal fits within a growing effort to build artificial intelligence virtual cells (AIVCs): models that learn to simulate cellular states from large-scale multi-omics measurements [<u>1</u>]. Recent advances in omics technologies, along with the release of open-source resources containing numerous experimental assays, support the scalable investigation of this problem. Datasets spanning different measurement technologies, gene spaces, and noise regimes offer a broad setting in which to ask whether methods successfully model cellular states and predict responses to perturbations. In particular, LINCS L1000 provides a diverse map of small-molecule perturbations with high-throughput expression assays of landmark genes, and Tahoe-100M provides a complete grid of single-cell chemical perturbation measurements over 50 cancer cell lines [<u>2</u>, <u>3</u>].

Predicting chemical perturbations is a notoriously hard modeling problem [4, 5]. Genetic perturbations often identify their direct molecular targets, thereby enabling models to use gene annotations, regulatory networks, or co-expression graphs as priors [6, 7]. In contrast, small molecules can have multiple targets [8], induce indirect downstream effects [9], with dose-dependent mechanisms [10]. Recent perturbation models attempt to capture this complexity using methods such as latent response shifts [11], disentangled covariate embeddings [12, 13], optimal transport maps [14], perturbation-conditioned generative models [15, 16], or attention-based gene–drug modelling [17]. Prior to the choice of modeling technique, a key challenge is defining how the perturbation is represented and provided as input, as model performance is highly sensitive to the chosen method [18]. Most strategies represent compounds using molecular fingerprints, chemical structures, or embeddings learned from structure by foundation models [19]. These natural choices are available before biological testing and can be directly computed for large compound libraries. Yet, chemical structure is a limited proxy for biological effect, as structural similarity is a weaker predictor of transcriptomic similarity than dose or cell line sensitivity [20].

In contrast, transcriptomic perturbation signatures provide rich information about the responses that compounds induce in specific cellular contexts. A single phenotypic readout captures on- and off-target effects, dose and exposure time, and activated downstream pathways. Building on this insight, we present BioPert, a simple framework for effective cross-context perturbation prediction based on the perturbation signature of a fixed reference cell line. We show that measured reference-cell transcriptomic responses serve as transferable perturbation representations and enable successful prediction of the full treatment-induced profile perturbation across cellular contexts, using a model architecture as simple as a multilayer perceptron. We compare this representation to an extensive array of molecular fingerprints and foundation model embeddings, and evaluate predictors on two chemical-perturbation resources, Tahoe-100M and LINCS L1000.

Across both datasets, chemical similarity, molecular fingerprints, and molecular foundation models show only limited practical gains over random controls. In contrast, BioPert predicts target cell line responses with a Pearson correlation of 0.79 on Tahoe-100M, a relative improvement of 75.5% over the next-best representation method. On LINCS, BioPert improves by 22.8% over the second-best method but only reaches an absolute Pearson correlation of 0.15. Further investigation demonstrates that this performance gap is linked to the variable reproducibility of profile replicates in LINCS, highlighting the necessity of mitigating batch effects of released resources. Crucially, BioPert far exceeds a strict baseline that copies the reference cell line response in the output, demonstrating that the model extrapolates to the target cell line context beyond simply reproducing reference-specific activity. Prediction performance remains robust across different choices of reference cell line, with only minor variation in aggregate performance. Although performance correlates with reference-target similarity, BioPert continues to predict target profiles with a Pearson correlation above 0.6 for the most dissimilar pairs, indicating that generalizability is not strongly dependent on a specific reference context.

Gene Set Enrichment Analysis (GSEA) [21] further confirms that BioPert predictions recover pathway-level activity. In an example of the melanoma line C32 exposed to cobimetinib, although pathways involved in MAPK-cascade signaling were activated in the reference, BioPert correctly predicted suppression in the target, indicating that the model captured compound-induced, cell-line-specific pathway activity. Overall, BioPert enables accurate chemical perturbation prediction by representing compounds in a form that captures their induced biological effects, even with a basic model architecture. However, these predictions come at the cost of experimental measurements, as they require access to high-quality transcriptomic perturbation signatures in a reference cell line for each compound of interest. Nevertheless, a single experimental readout allows extrapolation to multiple cellular contexts and could support collaborative virtual screening while maintaining the molecular composition of compounds private. We therefore envision that this representation method will serve as a valuable tool to reduce costs associated with phenotypic screening and to facilitate scalable drug repurposing.

## 2 Results

### 2.1 BioPert accurately predicts cross-context perturbation responses, while chemical fingerprints and learned embeddings offer limited gains over a random baseline

We trained simple multilayer perceptron (MLP) models to predict the transcriptomic responses, i.e., the Δ profiles, of target cell lines when exposed to treatments. Two distinct chemical perturbation datasets were considered: the single-cell dataset Tahoe-100M and the bulk, high-throughput transcriptomic profiling dataset LINCS L1000. The model input consisted of two concatenated vectors: one representing the target cell line’s identity, and one representing the treatment (see Figure 1). The target cell line was systematically represented by its average untreated control profile (DMSO), providing a measure of the cell line’s state in the absence of perturbation. In contrast, several options were explored for the treatment vector. The first combined a compound representation derived from its structure with dose and exposure-time information. We compared deterministic ECFP6 fingerprints [22] and embeddings from 14 foundation models based on molecular graphs, SMILES, or 3D geometry. These representations were scaled by treatment dose via a gating mechanism, and exposure time was appended as a one-hot encoding. The second option, a novel approach we named BioPert (*Bio*logical representation for *Pert*urbation prediction), was proposed as an orthogonal alternative to representations that reflect the molecular and structural features of the compounds. With BioPert, treatments were represented through the induced response (Δ profiles) of a fixed, held-out reference cell line (A549, unless otherwise stated) under matched treatment conditions. This representation does not consider the composition or arrangement of atoms and relies solely on the treatment’s effect on a reference cell line. Finally, random vectors were used as null controls, and each prediction was compared against two parameter-free baselines: the average of all training Δ profiles (*train* Δ) and direct copying of the reference cell line’s

**Figure 1:**
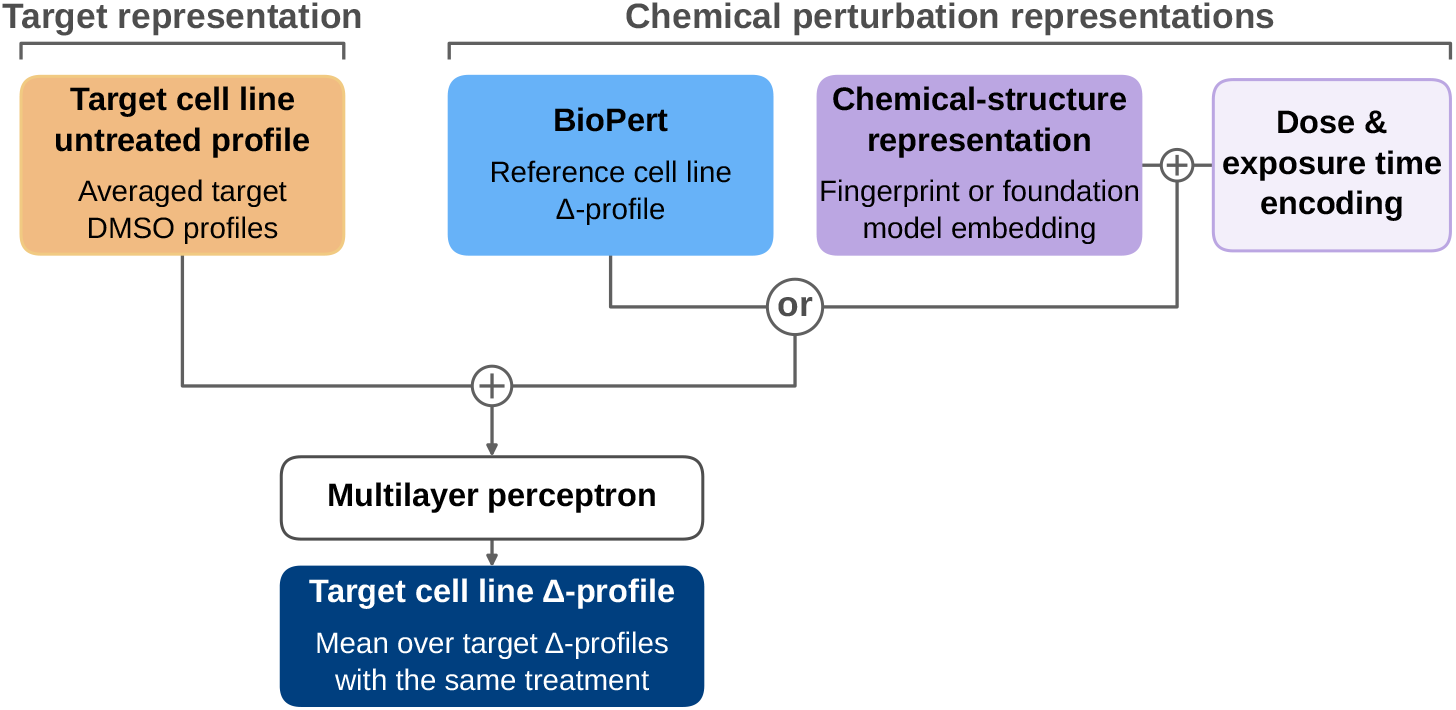
The model predicts a target cell line’s transcriptional response to a chemical perturbation from representations of the target cell line and treatment. The model concatenates the untreated target cell line profile with either BioPert (the treated-minus-untreated, i.e., Δ, profile in a reference cell line, which inherently captures dose and exposure time) or a chemical-structure representation (a fingerprint or molecular foundation-model embedding) paired with a separate dose and exposure-time encoding. An MLP maps these inputs to the predicted Δ profile in the target cell line.

Δ profile under matched treatments (*reference* Δ). All predictors were evaluated on the same test observations, with compounds held out from training, and allocated the same budget over a random search on hyperparameters, including model architecture and optional input dimensionality reduction.

Chemical representations yielded only marginal improvements over random vectors in both Tahoe-100M and LINCS (Figure 2d-e). In Tahoe-100M, the best structure-derived encoder, ChemBERTa-77M-MLM, achieved an average test Pearson correlation of 0.451 (95% CI, 0.447–0.454), a relative improvement of 2.8% over the random-vector control (0.439; 95% CI, 0.434–0.442). Morgan fingerprints and the other molecular embeddings performed similarly to the random control, indicating that the chemical geometry encoded by these representations contributed little predictive signal. The same pattern held in LINCS, although absolute correlations were lower: the best encoder, CheMeleon, achieved an average test Pearson correlation of 0.123 (95% CI, 0.122–0.125), a 12.9% improvement over the random-vector control (0.109; 95% CI, 0.108–0.111). Together, these results indicate that the mapping from structure to transcriptomic response is not captured by a simple monotonic relationship in the evaluated representation spaces and is difficult to learn from the available data.

**Figure 2:**
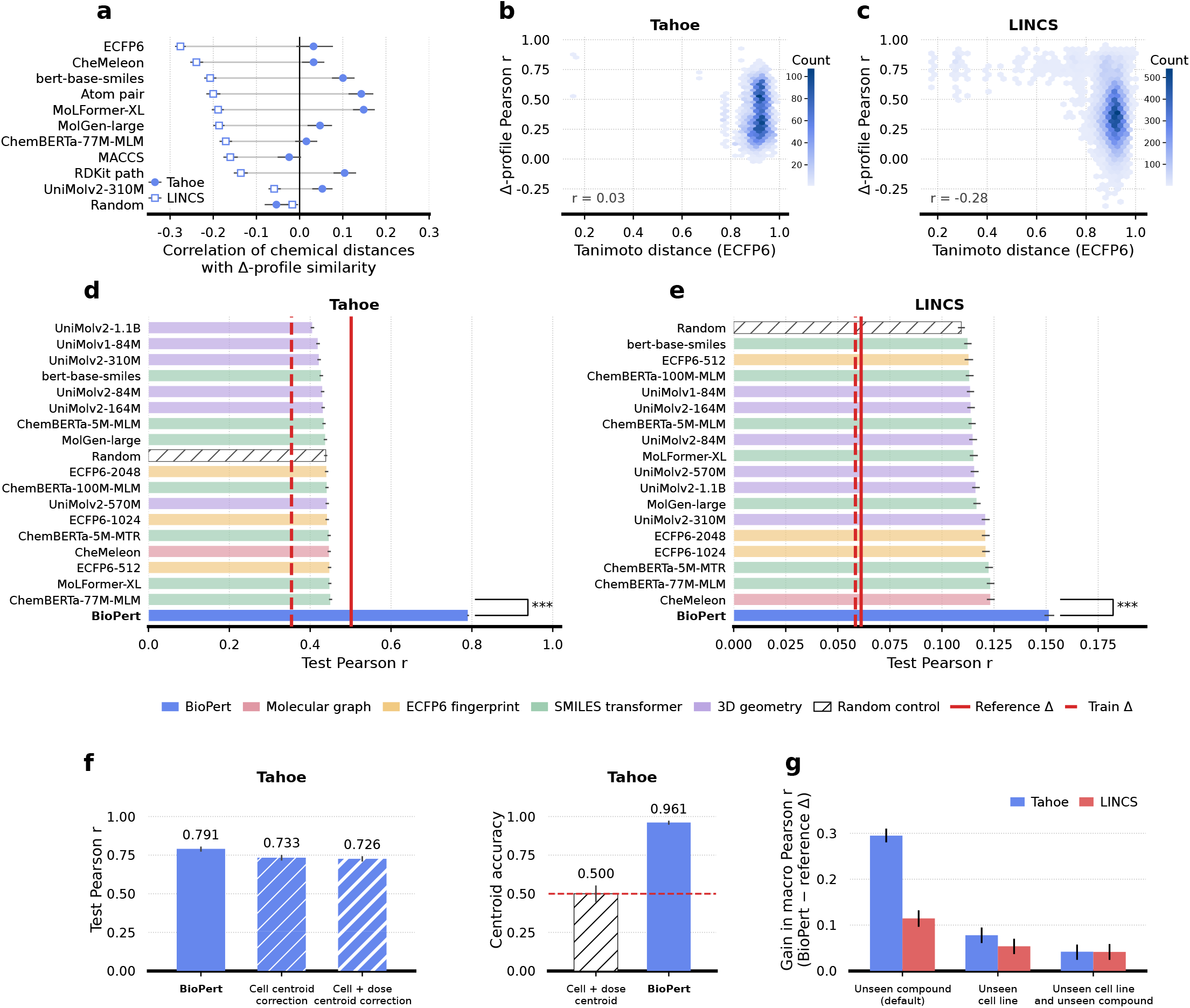
Molecular fingerprints and foundation-model representations provide limited gains over a random control, whereas BioPert allows cross-context perturbation effect prediction. **a**. Pearson correlation between molecular distances (Tanimoto or cosine) and transcriptomic Δ-profile similarity (Pearson correlation) of pairs of compounds. Filled circles show Tahoe-100M, open squares show LINCS full results are available in Supplementary Fig. 4). **b**,**c**. Pairwise Δ-profile correlation as a function of pairwise ECFP6 Tanimoto distance between compounds in Tahoe-100M (b) and LINCS L1000 (c). Hexagon color denotes the number of compound pairs. **d**,**e**. Average test Pearson correlation between true and predicted Δ profiles for held-out compounds, for predictors trained with varying treatment representations on Tahoe-100M (d) and LINCS L1000 (e). Color encodes representation type (Morgan fingerprint, SMILES transformer, 3D geometry, molecular-graph MPNN, random control, and BioPert). Two parameter-free baselines are drawn as vertical lines (*reference* Δ and *train* Δ). **f**. Tahoe Systema sensitivity analysis: Pearson correlation before and after subtracting training-derived generic perturbation centroids (left), and Systema centroid accuracy against other drugs measured in the same target cell line and dose (right). BioPert’s performance is robust to corrections accounting for systematic effects. **g**. BioPert absolute improvements over the *reference* Δ baseline in different generalization settings. While gains over the baseline are substantial for held-out compounds, predictions in novel cellular contexts are more challenging.

In contrast, BioPert achieved an average test Pearson correlation of 0.791 on Tahoe-100M (95% CI, 0.788–0.793) and 0.151 on LINCS (95% CI, 0.149–0.154). The performance differences between LINCS and Tahoe-100M are explained in section 2.2. Relative to the best chemical representation, BioPert improved by 75.5% on Tahoe-100M and 22.8% on LINCS. BioPert also significantly outperformed both parameter-free baselines in Tahoe-100M (*reference* Δ, *P <* 0.001; *train* Δ, *P <* 0.001) and LINCS (*reference* Δ, *P <* 0.001; *train* Δ, *P <* 0.001). On Tahoe, it improved over the *reference* Δ baseline by 57.7%, demonstrating that the model extrapolated beyond reproducing the reference response and adapted predictions to the context of the target cell line. Restricting Tahoe-100M predictions to 964 LINCS landmark genes produced a similar average correlation (0.792; 95% CI, 0.790–0.794), indicating that the performance was not attributable to gene-set size. On LINCS, BioPert achieved higher accuracy with less training data than the chemical encoders (225,562 vs. 369,199), as it required that each compound be profiled in the reference cell line. Notably, this representation method does not require encoding additional conditions, such as dose and exposure time, because these variables are implicitly captured in the reference responses. Nevertheless, it requires experimentally profiling the reference’s response for each compound, thereby trading off additional experimental cost for improved predictions.

Because correlations computed relative to untreated controls can be inflated by responses shared across perturbations, we next evaluated BioPert using the Systema framework [23]. In Tahoe-100M, generic perturbation centroids were estimated exclusively from training compounds, either separately for each target cell line or for each combination of target cell line and dose, and subtracted from both observed and predicted Δ profiles. BioPert retained most of its predictive power after this correction: the average Pearson correlation decreased from 0.791 without centering to 0.733 (95% CI, 0.715–0.752) after cell-line-specific centering and 0.726 (95% CI, 0.708–0.743) after the stricter cell-line and dose centering (Figure 2f). Moreover, BioPert predictions were closer to their matched compound response than to 96.1% of alternative compound responses measured in the same target cell line and dose (95% CI, 94.7–97.3%), compared with a 50.0% baseline for the centroid perturbation. While previous experiments tested predictors on held-out compounds, we further assessed generalization across cellular and chemical contexts by holding out specific cell lines from the training set. BioPert achieved smaller gains over the *reference* Δ baseline for held-out cell lines exposed to previously observed compounds (Tahoe-100M: 0.078, 95% CI, 0.061–0.095; LINCS: 0.054, 95% CI, 0.037–0.070) and for the strict intersection of independently held-out cell lines and compounds (Tahoe-100M: 0.042, 95% CI, 0.024–0.058; LINCS: 0.042, 95% CI, 0.024–0.059; Figure 2g). Thus, although BioPert still learned context-dependent transformations that produced gains over the *reference* Δ baseline, predictions in held-out cellular contexts were more challenging.

### 2.2 Prediction performance is associated with measurement quality and reproducibility of condition replicates

Across representation methods, performance on LINCS was substantially lower than on Tahoe-100M. We found that this gap likely reflects batch effects and the limited reproducibility of perturbation responses in LINCS. When comparing pairs of samples replicated in identical conditions, absolute profiles were individually highly reproducible, with a median pairwise Pearson correlation of 0.92 across intra- and inter-plate comparisons (Figure 3a, absolute profiles). However, reproducibility declined sharply when isolating the perturbation effect by considering Δ profiles. The median correlation of matched Δ profiles within the same plate (intra) was measured at 0.45, and dropped to 0.07 when considering samples from different plates (inter). In addition, intra-plate reproducibility was primarily driven by the 20 *µ*M, 24 h condition, which yielded the largest mean response magnitude (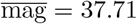, *n* = 35,476) and a median replicate correlation of 0.75, compared with 0.28 for all other dose and time combinations (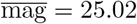*n* = 79,494; Figure 3a, intra-plate).

**Figure 3:**
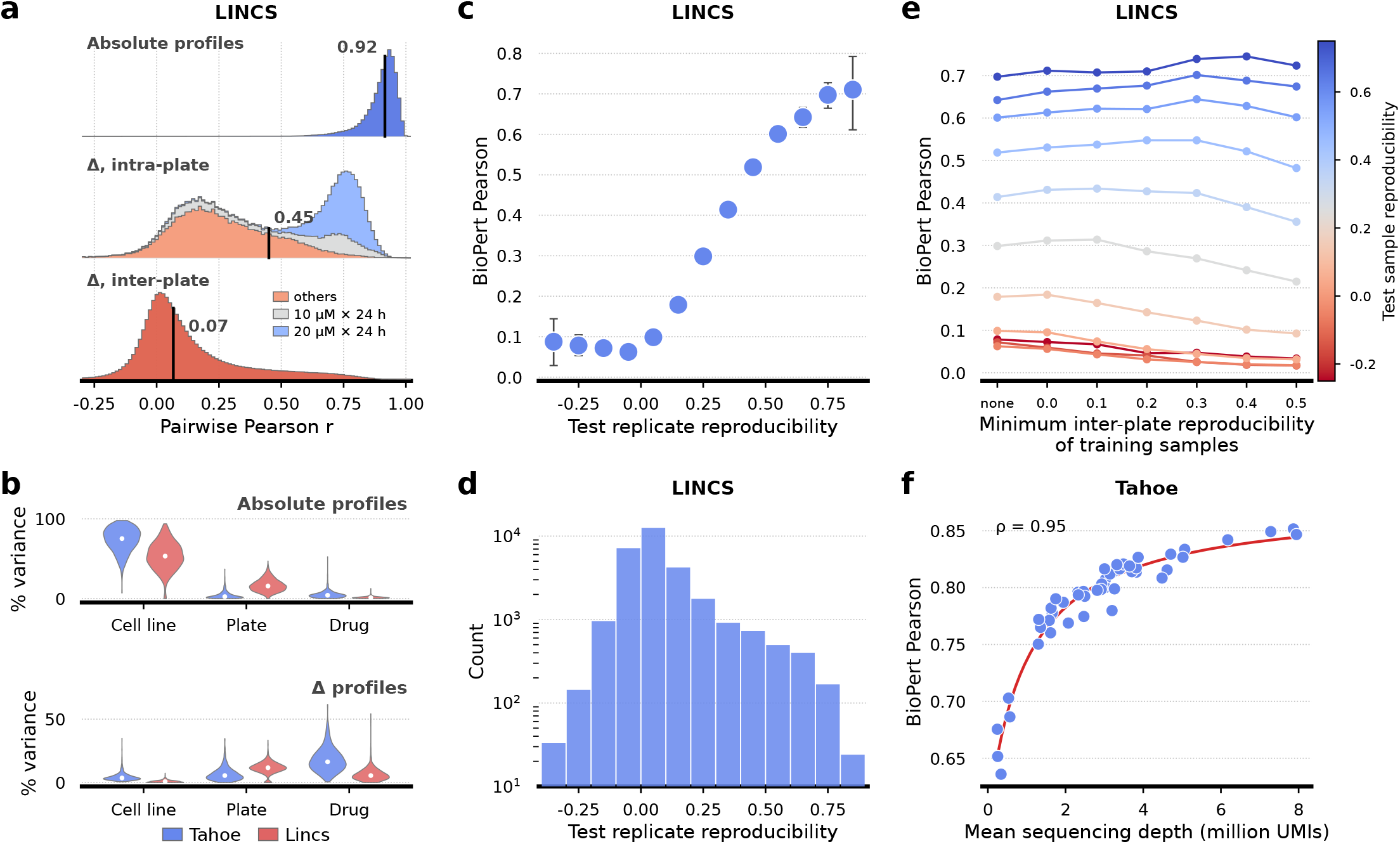
Experimental measurement quality and batch effects constrain achievable performance. **a**. Distributions of Pearson correlation between pairs of replicate conditions on LINCS, for absolute and Δ profiles, computed separately for intra-plate and inter-plate replicate pairs. Vertical lines mark the median of each distribution. The bimodality of the intra-plate distribution is accounted for by the perturbation conditions: replicates exposed to the highest dose (20 *µ*M for 24 h) show higher reproducibility than all other conditions. **b**. Variance decomposition analysis (lme4 REML), reporting the share of expression variance attributable to experimental factors for absolute profiles (top) and Δ profiles (bottom) in both datasets. Plate identity always accounts for a higher proportion of the expression variance in LINCS than in Tahoe. **c**. BioPert performance on the test set as a function of the reproducibility of the test conditions (inter-plate, binned). Prediction correlations with the targets are monotonic in target reproducibility. **d**. Number of test conditions per reproducibility bin (log scale).**e**. Test Pearson on LINCS as a minimum reproducibility threshold for the training set is increased from none to 0.5, with the test set split into fixed reproducibility bins. High-reproducibility bins slightly increase while low-reproducibility bins degrade as more training data is discarded. **f**. Average test Pearson per cell line, versus mean sequencing depth of the pseudobulk samples (UMIs per condition), on Tahoe. Performance increases with sequencing depth.

This unfavorable signal-to-noise ratio was confirmed by running an independent expression variance decomposition analysis using the *variancePartition* framework [24]. On Tahoe, the per-gene median expression variance attributable to plate identity was 2.4% for absolute profiles and 5.5% for Δ profiles. In contrast, plate identity accounted for 15.7% and 11.7% of the median gene-level variance in absolute and Δ profiles, respectively, in LINCS. This confirms the presence of expected batch effects, given LINCS is a collection of high-throughput assays conducted in variable experimental settings.

This lack of measurement reproducibility is associated with low prediction performance. After binning test samples by reproducibility levels, correlations of BioPert predicted profiles with the ground truth increased monotonically with test-condition reproducibility (*R*^2^ = 0.92 for the binned means; condition-level Spearman *ρ* = 0.39, 95% CI, 0.38–0.40, *P <* 0.001; Figure 3c). In particular, prediction correlations approached 0.7 for the most reproducible LINCS bins, comparable to BioPert’s performance on Tahoe-100M. While most Tahoe-100M conditions lack replicates, which prevents comparable reproducibility analyses, we used sequencing depth (UMI count) as a proxy for the reliability of target measurements. Average prediction correlation increased with the mean UMI count of pseudobulk profiles across cell lines (Spearman *ρ* = 0.95, 95% CI, 0.88–0.98, *P <* 0.001; Figure 3f), further indicating that the quality of the experimental measures constrained prediction performance.

This analysis also prompted us to examine whether excluding poorly reproducible conditions from the training set improved predictions. We set a minimum reproducibility threshold and filtered LINCS training conditions accordingly. Increasing the threshold slightly improved performance on highly reproducible test samples but reduced accuracy on poorly reproducible ones. As the threshold increased from no filtering to a minimum correlation between replicates of 0.5, the overall average test correlation decreased from 0.151 to 0.084, while the training set was reduced from 225,562 to 7,492 samples (Figure 3e). Because filtering removed more than 96% of the training data, gains in training-set reproducibility were traded against a loss of perturbation coverage.

### 2.3 Biological-response representations are robust to the choice of reference cell line and improve when more replicates are available

The selection of the reference cell line had only a modest influence on the average predictive performance. However, greater similarity between the reference and target cell lines was associated with improved performance for specific targets. Replacing the A549 reference with three alternative cell lines resulted in a shift of less than 0.03 in the average test Pearson correlation on Tahoe-100M (ranging from 0.785 to 0.810 across A549, A-172, COLO 205, and PANC-1) and a comparable shift on LINCS (from 0.135 to 0.151 across A549, MCF7, and PC3) (Figure 4a,b). These findings indicate that biological-response representations are not dependent on the unique characteristics of a single reference cell line. However, on Tahoe, a positive correlation was observed between the target-reference response similarity and the average performance on the target (Spearman *ρ* = 0.79, 95% CI, 0.72–0.85, *P <* 0.001). This indicates that reference responses close to target responses support improved predictions. Nevertheless, even the lowest reference–target response similarities, below 0.25 in Pearson correlation, enabled predictions with correlations above 0.60 to ground-truth responses.

**Figure 4:**
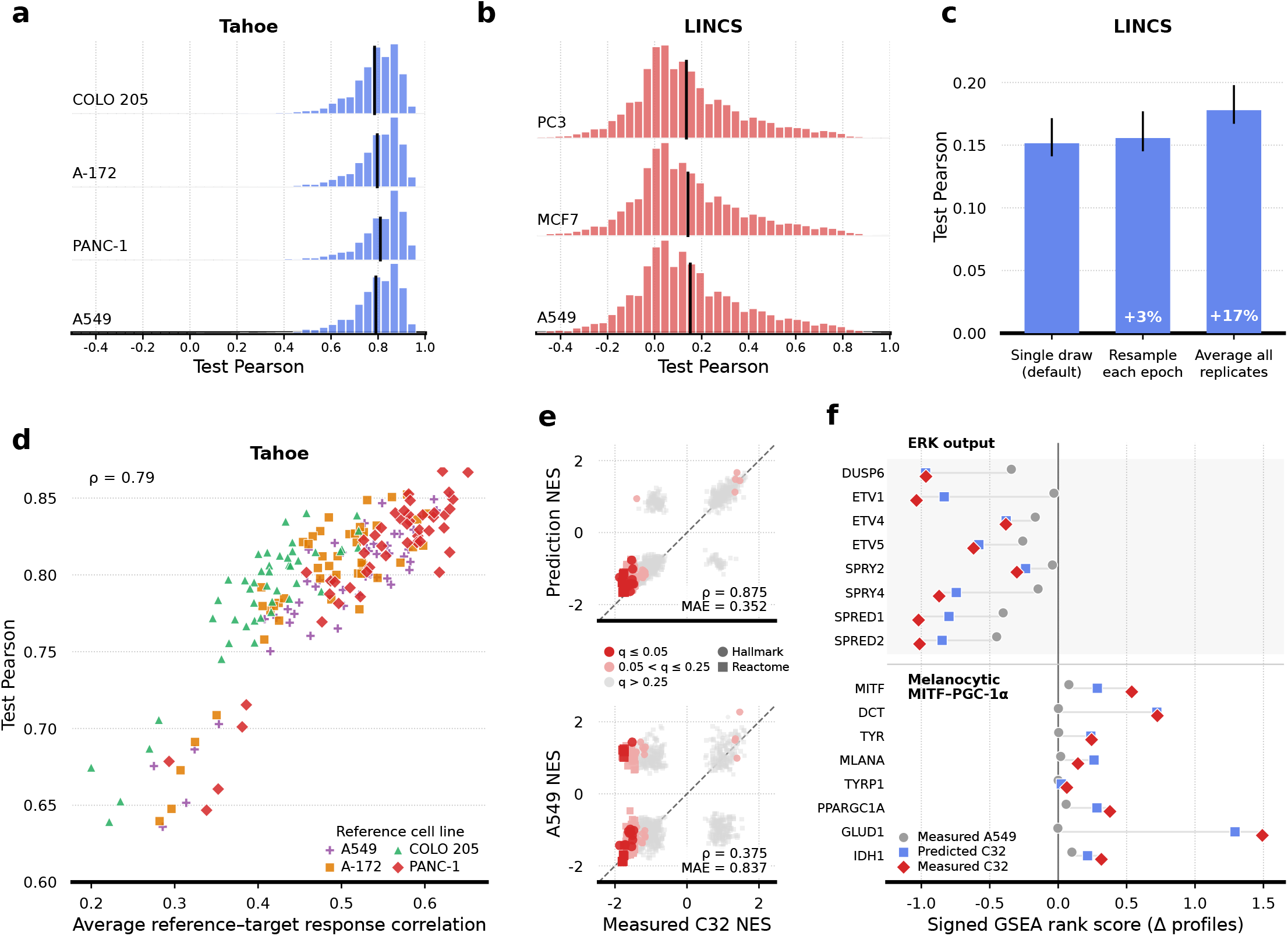
BioPert representations are robust across reference cell line choices and support predictions in target contexts differing from the reference. **a**,**b**. Distribution of mean test Pearson correlation for each target cell line under different choices of reference cell line, on Tahoe-100M (a) and LINCS L1000 (b). **c**. Test performance on LINCS when varying the replicate aggregation scheme of input reference Δ profiles. Re-sampling among replicates at each epoch improves performance over selecting a fixed representative, and averaging all replicates provides further gains. **d**. Test performance for each cell line versus the similarity between the reference and target cell’s Δ profiles, for all four reference cell lines. Performance increases with reference–target similarity. **e**. For cobimetinib (0.5 *µ*M, 24 h) in the BRAF-V600E melanoma line C32, measured target pathway NES versus either BioPert-predicted C32 NES or measured A549 reference NES across Hallmark and Reactome pathways. **f**. Signed ranking scores for established ERK-output genes and a melanocytic MITF–PGC-1*α* module in measured C32, predicted C32, and measured A549. BioPert successfully recovers target-dependent pathway activations that differ from the reference.

Moreover, on LINCS, enhancing the quality of the reference signature improved predictive performance. The default configuration employed a single replicate, fixed throughout training, to mirror the realistic scenario in which each compound is assayed once in a reference cell line. When additional replicates were available, resampling them across training epochs led to performance gains, while averaging replicates yielded the highest performance (+17.4% relative increase; Figure 4c). The averaging process reduced replicate-specific noise and provided a more accurate estimate of the shared compound response. In contrast, resampling introduced replicate variability but retained noise in each input. Therefore, repetitive replicate profiling can directly improve prediction accuracy, although this approach comes at higher experimental costs.

### 2.4 Predicted profiles recover a melanoma lineage-specific response to MEK inhibition

To assess whether BioPert’s predictions reflect target-specific biology rather than reference-line inheritance, we examined cobimetinib (a MEK1/2 inhibitor) response in the BRAF-V600E melanoma line C32, predicted from the KRAS-mutant lung adenocarcinoma reference A549. This example was chosen for its maximal divergence in both oncogenic driver and cell lineage. Across the merged set of Hallmark and Reactome pathways, predicted C32 pathway activity closely matched the measured target response (Figure 4e: Spearman *ρ* = 0.875), whereas the measured A549 reference diverged substantially (*ρ* = 0.375). This includes a subset of pathways centered on MAPK-cascade signaling (MAP2K/MAPK activation, signaling by high-kinase-activity BRAF mutants) that were suppressed in the target but activated in the reference, with the prediction correctly following the target. This divergence is consistent with the differential dependence of BRAF-versus KRAS-mutant lines on sustained MAPK output [25, 26], and with reported adaptive pathway reactivation in KRAS-mutant models upon MEK inhibition [27, 28].

Moreover, at the gene level, we examined canonical ERK-output genes (DUSP6, ETV1, SPRY4, SPRED1/2), transcriptional readouts of ERK activity that serve as direct pharmacodynamic markers of MEK inhibition. These genes were coordinately suppressed in both measured and predicted C32 profiles but largely unchanged in A549 (Figure 4f). Reciprocally, a curated MITF–PGC-1*α* module [29, 30] was induced in measured and predicted C32 but flat in the non-melanocytic reference. This module comprises melanocytic differentiation genes (MITF, DCT, TYR, TYRP1, MLANA) and oxidative-phosphorylation genes (PPARGC1A and downstream TCA-cycle targets). Although C32 is amelanotic, it retains this transcriptional machinery in a suppressed state at baseline. MEK inhibition relieves this suppression, consistent with prior reports in BRAF-mutant melanoma [31, 29]. Because this lineage-specific program has no counterpart in A549’s biology, its correct retrieval indicates that the prediction reflects the target’s specific biology rather than the reference’s post-treatment expression profile. Altogether, these results highlight that BioPert’s predictions track target-specific, genotype- and lineage-dependent transcriptional responses to cobimetinib.

## 3 Discussion

Our results show that a compound’s measured transcriptional response is a more informative representation of its biological activity than its chemical structure alone. Across Tahoe-100M and LINCS L1000, molecular fingerprints and foundation model embeddings provide only limited gains over random compound representations, whereas representations based on a reference cell line’s responses substantially improve predictions. This advantage persists across single-cell and bulk measurements, different gene spaces, distinct noise regimes, and experimental technologies. On Tahoe-100M, BioPert predicts responses that, on average, correlate with the ground truth at 0.791. This result is achieved with a simple MLP, underscoring the effectiveness of designing task-appropriate representations. Moreover, BioPert implicitly encodes complete experimental conditions such as dose and exposure time without requiring additional design choices.

Notably, these results are achieved on predictions for the isolated treatment-induced response rather than the absolute treated state. Several perturbation-prediction frameworks evaluate predictions in the expression space of treated cells [11, 12, 13]. Because basal cell identity dominates both untreated and treated profiles, high correlations between absolute profiles can largely reflect reconstruction of the target cell line rather than prediction of the perturbation effect. Consistent with this concern, untreated and treated replicates are highly correlated in both datasets, while replicate Δ profiles are substantially less reproducible. Plate-matched baseline subtraction therefore makes the prediction task more difficult, but it also aligns the evaluation with the biological quantity of interest: the transcriptional change attributable to treatment.

Moreover, model abilities directly rely on experimental measurement quality. Performance increases with the sequencing depth of test conditions on Tahoe, and the lower accuracy on LINCS is associated with poor inter-plate reproducibility of test samples. While simple filtering of noisy training conditions does not improve measurable performance, improving the reference signature itself is beneficial. Our default setting uses one fixed replicate throughout training, reflecting the realistic experimental scenario in which each compound is assayed only once in a reference cell line. When additional replicates are available, resampling them across training epochs improves performance, and averaging them performs best. Replicate profiling is therefore a direct route to better predictions, but the gain comes with increased experimental cost. Finally, changing the reference cell line has a modest effect, and cross-context predictions remain accurate even when target responses differ from the reference, indicating that this method does not rely on a uniquely privileged reference context.

A possible experimental workflow could involve a conceptual separation between compound characterization and assay calibration. Transcriptomic profiling can define a reusable biological coordinate system for compounds, onto which additional cellular contexts are mapped using comparatively limited target-specific information. Determining how many compounds are required to calibrate a new assay, and whether multiple reference cell lines provide complementary information beyond a single well-chosen reference, could further help assess the cost–accuracy trade-off of this strategy.

## 4 Methods

### 4.1 Definitions

We refer to a chemical perturbation induced by a compound at a given dose and exposure time as a *treatment*. Its treatment-induced transcriptional response in a cell line is termed the Δ profile (see Section 4.2.3). We call the cell line for which the perturbation is predicted the *target cell line*, and the cell line whose Δ profile is used to represent a treatment the *reference cell line* (see Section 4.3.1).

### 4.2 Data preprocessing

#### 4.2.1 Tahoe-100M

We aggregated raw single-cell RNA-seq data from Tahoe-100M [3] into pseudobulks, one per unique (cell line, treatment well) pair, where each well encodes a single drug–dose combination applied for 24 h. For each pseudobulk, raw UMI counts were summed across all constituent cells and restricted to 19,020 protein-coding genes as defined by the HGNC reference [32]. Pseudobulks with fewer than 50 cells or fewer than 35,000 UMIs among these protein-coding genes were discarded, and cell lines with more than 90% discarded pseudobulks were entirely removed. Expression values were log-normalized as

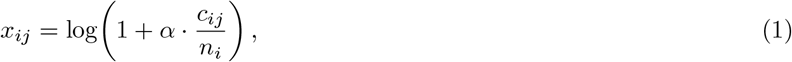

where *c*_*ij*_ is the raw UMI count for gene *j* in pseudobulk *i, n*_*i*_ is the total UMI count of pseudobulk *i*, and *α* = 10,000. Throughout this work, we refer to these log-normalized pseudobulks as Tahoe-100M profiles. The two SMILES [33] missing from the dataset metadata were retrieved from PubChem (Sacubitril/Valsartan, PubChem CID 24755620 and Verteporfin, PubChem CID 168430535).

#### 4.2.2 LINCS L1000

LINCS L1000 profiles [2] were downloaded from the CLUE data dashboard (https://clue.io/releases/data-dashboard) as the Phase II (LINCS 2020 beta) release on 2025-12-28. We restricted the data to Level 3 (quantile-normalized) profiles and retained only the 978 directly measured landmark genes. We further filtered profiles to DMSO vehicle controls `(pert_type = ctl_vehicle)` and small-molecule-treated measurements `(pert_type = trt_cp)`, keeping only those passing quality control `(qc_pass = 1).` We then removed treated profiles lacking a matching DMSO control, where a match required the same cell line, and detection plate. Finally, we discarded profiles with missing values in any required metadata field (cell line, perturbagen identifier, SMILES, dose, exposure time, detection plate, or sample identifier).

#### 4.2.3 Treatment response (Δ profiles)

We isolated the treatment-induced transcriptional response from the cell-line-specific basal expression that otherwise dominates a treated profile. To this end, for each treated profile **x**_trt_, we computed a corresponding Δ profile as the gene-wise difference from the mean of matched DMSO controls, where matching was performed on cell line, and plate within the same dataset:

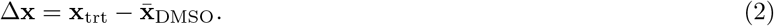

### 4.3 Treatment representation

#### 4.3.1 Biological perturbation signature

##### BioPert

Treatments were represented by the induced Δ profiles in a fixed reference cell line. One profile was randomly selected for each target observation and held fixed throughout training and evaluation. This design approximates the common experimental setting in which each treatment is measured once in a reference cell line. As a controlled ablation, this fixed single-profile strategy was compared with resampling among available reference replicates at each training epoch and with averaging all available reference replicates (Figure 4c). Per-epoch resampling acted as data augmentation by introducing replicate variability into the treatment representation, while averaging reduced replicate-specific noise in the reference representation.

##### Reference cell line

Unless otherwise specified, A549 was used as the default reference cell line and was excluded from the prediction targets in these experiments. A549 is a human lung cancer cell line [34] widely used as a model in drug screening and preclinical studies [35]. It was initially selected as it is common to both Tahoe-100M and LINCS and is the most-represented cell line in LINCS after pre-processing.

In Tahoe, the number of treated profiles is comparable across cell lines, a consequence of the cell village protocol, in which all cell lines are pooled and treated together within a single well [3]. In LINCS, however, the number of treated profiles per cell line varies widely, constraining the available data by requiring that the reference and target profiles be mapped under the same treatment conditions. In LINCS, using A549 as the reference cell line reduced the number of doses from 65 to 52, spanning 0.0001–200 *µ*M ( *≈* 6.3 orders of magnitude), and reduced the number of exposure times from 11 to 4, ranging from 3 h to 48 h. Nevertheless, subsequent ablations with other reference cell lines demonstrated that this choice had limited impact on the downstream performance (Figure 4a,b).

#### 4.3.2 Molecular representations

BioPert was compared against a panel of molecular representations spanning several families and a control baseline:

1. Molecular fingerprints: Extended Connectivity Fingerprints with radius 3 (ECFP6) [22] at three bit-vector lengths (512, 1024, and 2048), computed from SMILES using RDKit’s [36] `rdFingerprintGenerator.GetMorganGenerator.`
2. Pretrained molecular embeddings from foundation models:
  a. String-based transformers: ChemBERTa-5M-MLM, ChemBERTa-5M-MTR, ChemBERTa-77M-MLM, and ChemBERTa-100M-MLM [37]; MoLFormer-XL-both-10pct [38]; `unikei/bert-base-smiles` [39]; and MolGen-large [40]. For each model, we used `pooler_output` when exposed and otherwise mean-pooled the final hidden layer. MolGen inputs were converted from stripped SMILES to SELFIES [41].
  b. 3D-geometry-based models: Uni-Mol v1 84M [42] and Uni-Mol2 84M, 164M, 310M, 570M, and 1.1B [43], generated from stripped SMILES using `unimol_tools.UniMolRepr.`
  c. Molecular-graph model: CheMeleon [44], using the fingerprint extracted by the released pretrained model.
3. Random control: one fixed 512-dimensional vector of independent standard-normal values per exact SMILES, independent of molecular structure.

##### Dose and time encoding

Directly comparing BioPert to molecular representations would confound predictions with a genuine difference in information content: a Δ profile implicitly encodes the dose and exposure time of the treatment, whereas a molecular representation depends on chemical structure alone and is invariant to the conditions under which the compound is applied. This confound was removed by explicitly conditioning the predictions on treatment dose and time. Three different encoding schemes were assessed:

1. Dose as a scalar concatenated to the model input: log_10_(*d*), where *d* is the dose in *µ*M.
2. Dose binned into categorical bins: for Tahoe, one bin per each of the 3 discrete dose values; for LINCS, seven bins with boundaries fixed *a priori* at each power of ten, covering the full dose range (*≈*6.3 orders of magnitude).
3. A multiplicative doser gate, adapted from the dose encoder in the CPA [12] codebase^1^, in which the embedding of the compound is scaled by:

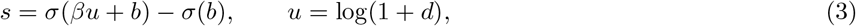

where *σ*( *·* ) denotes the sigmoid function, *d* is the dose in *µ*M, and *β, b* are learned scalar parameters. (*β, b*) were initialized to (1, 0), shared across compounds, and learned jointly by backpropagation.

The three candidate encodings were compared against a blind control, in which the model received no dose information, for four compound representations: random vectors (*n*_dim_ = 512), ECFP6 [22] (Morgan fingerprint [45]) (*n*_dim_ = 1024), CheMeleon [44], and Uni-Mol2 1.1B [43]. The multiplicative gate performed best in validation performance in all cases, and we therefore adopted this dose encoding for all subsequent experiments using molecular representations (Supplementary Fig. 6). Finally, exposure time was one-hot encoded for LINCS and omitted for Tahoe-100M, which includes only the 24-hour setting.

### 4.4 Structure–activity relationship analysis

To assess whether transcriptomic responses reflect compound similarity, we compared 25 representations: 10 molecular fingerprints, 14 foundation-model embeddings, and the random-vector control. Fingerprints were compared using Tanimoto distance, whereas continuous embeddings and random vectors were compared using cosine distance.

For fingerprints, the symmetric Tanimoto similarity matrix *T* ***∈*** [0, 1]^*n×n*^ was constructed as

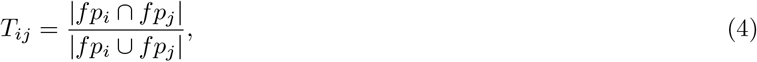

where *fp*_*i*_ denotes the bit-vector fingerprint of compound *i*. The corresponding Tanimoto distance was *D*_*ij*_ = 1 *− T*_*ij*_. For foundation-model embeddings, we computed cosine distance between each pair of embedding vectors. We first retained conditions with a defined mean inter-plate Δ-profile correlation of at least 0.7. Within each shared assay, one randomly drawn replicate was retained per SMILES. We then formed every unordered compound pair in the assay. Transcriptomic similarity for a pair was the Pearson correlation between its two Δ profiles, and profile magnitude was the *ℓ*_2_ norm of each profile. For each representation, we calculated the Pearson association between representation distance and transcriptomic similarity across all eligible assay-specific pair rows.

For both Tahoe-100M and LINCS, there was weak to no association between ECFP6 Tanimoto distance and output Δ profile similarity (Pearson *r* = 0.03 and *−* 0.28, respectively; Figure 2b-c). This finding was not attributable to limited structural diversity of the compounds, as they had mean pairwise distances of 0.92 in Tahoe-100M and 0.91 in LINCS. This weak association persisted across the total of 25 molecular representations evaluated: correlations with paired Δ profile similarities remained below 0.4 in magnitude, generally below 0.2, varied in direction between datasets, and were not stronger for pretrained embeddings than for simple fingerprints (Figure 2a).

### 4.5 Model and training

#### Model

A simple MLP was implemented in Julia using the Flux.jl library [46]. As input, it received concatenated representations of a target cell line and a treatment. The target cell line was represented by the unweighted mean of all its DMSO profiles, and treatment representation methods are introduced in subsection 4.3. Optional dimensionality reductions were applied to both representations. In the output, target Δ profiles over the complete gene space with matched compound, dose, and exposure time were averaged across plates using an unweighted arithmetic mean.

#### Input dimensionality reduction

Principal Component Analysis (PCA) was applied independently to the treatment and target-cell input representations. Each PCA was fit to the full training-observation matrix and then applied to the validation and test observations. When BioPert was used to represent the treatment, the number of retained principal components (PCs) was the same for both the Δ profile of the reference cell line and the untreated profile of the target cell line: *n*_PCA,expr_. If per-epoch resampling of the reference cell line’s Δ profile was used, the PCA model was not refit on the resampled replicates. When the model used a molecular embedding to represent the treatment, dose scaling and concatenation with the exposure-time one-hot vector were performed after dimensionality reduction.

#### Landmark-gene experiment

In an additional experiment, Tahoe-100M profiles were restricted to the 964 genes intersecting with the 978 genes measured directly by LINCS L1000. A separate BioPert model was trained and tuned on this restricted gene space using the same compound split and selection procedure as the main Tahoe-100M model.

#### Baselines

The *train* Δ baseline is the global gene-wise mean of all training-observation Δ profiles. The *reference* Δ baseline is the unweighted mean of all Δ profiles of the fixed reference cell, matching the target compound, dose, and exposure time.

#### Data split

To assess generalization to structurally distinct held-out compounds, we constructed train/validation/test splits by clustering compounds on chemical structure using Butina clustering [47]:

1. An ECFP6 fingerprint (2048 bits) was computed for each unique SMILES using RDKit’s [36] `GetMorganGenerator.`
2. Pairwise Tanimoto similarities between ECFP6 fingerprints were computed using RDKit’s `BulkTanimotoSimilarity` and converted to a distance matrix as one minus the similarity.
3. Compounds were partitioned into clusters using RDKit’s `Butina.ClusterData` with a Tanimoto distance cutoff of 0.4. The algorithm returns clusters ordered by decreasing size.
4. Train, validation, and test fractions were set to 0.75, 0.10, and 0.15, respectively. Clusters were assigned greedily, first filling the training set until it reached its target fraction, then the validation set, with any remaining clusters assigned to the test set.

Representation comparisons used the common intersection of eligible test observations. The resulting split was fixed and reused as the default setting across all comparisons. As a result, the training, validation, and test set comprised 283, 38, and 56 compounds in Tahoe (39,685, 5,260, and 7,767 observations), and 5,787, 772, and 1,157 compounds in LINCS (225,562, 28,630, and 39,980 observations).

#### Cell line generalization

Cell lines were randomly assigned to independent, fixed training, validation, and test sets in addition to the compound split. Two settings were evaluated in Fig. 2g. In the observed-compound setting with held-out cell lines, test compounds were seen during training in other cell lines, while target cell lines were absent from training. In the joint setting, both the compound and target cell line belonged to their independently defined test sets. The full-dataset DMSO mean of each held-out target cell line and its matched A549 responses remained available as model inputs. Each generalization configuration was tuned independently. Performance gain was calculated per test observation as the prediction Pearson correlation minus the correlation obtained by the *reference* Δ baseline, and was then averaged within target cell lines.

#### Training

Fully connected hidden layers used ReLU activations, and the output layer was linear. We used RMSE as the training loss function. Models were trained separately on Tahoe and LINCS using Flux.jl [46] and AdamW [48] (*β*_1_ = 0.9, *β*_2_ = 0.999). The learning-rate schedule linearly increased from 10^*−*8^ to the trial learning rate over the first ten epochs and then used cosine annealing [49] to one-tenth of that rate. All non-bootstrap stochastic procedures, including preprocessing, splitting, initialization, fixed reference selection, and training, used seed 42.

#### Hyperparameter search

Hyperparameter configurations were sampled by random search using Weights & Biases [50] sweeps. Each combination of dataset and treatment representation received 50 trials with an identical search space, which included MLP architecture and input PCA dimensions. Notably, dimensionality reduction was optional for molecular representation methods. Detailed ranges for each hyperparameter are provided in Supplementary Methods. We checkpointed Pearson and Spearman correlations every ten epochs and performed best-trial and best-checkpoint selection based on average validation Spearman correlation. We report the corresponding average Pearson correlation on the test set as model performance.

#### Statistical analysis

We estimated confidence intervals by running bootstrap analyses with 1,000 resamples, seed 0, and 95% percentile confidence intervals. Main pairwise model comparisons used two-sided paired Wilcoxon signed-rank tests on matched per-observation correlations. Benjamini–Hochberg correction was applied to the Wilcoxon tests in Fig. 2d,e, with Tahoe-100M and LINCS as separate families, and to the paired comparisons across test-reproducibility bins in Fig. 3e.

### 4.6 Systema analysis

We used the Systema framework [23] to test whether predictions captured compound-specific responses beyond generic perturbation effects. From training observations only, we calculated a centroid for each target cell line *l* and for each context defined by target cell line and dose (*l, d*):

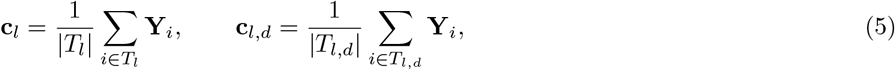

where *T*_*l*_ and *T*_*l,d*_ contain the corresponding training observations. The relevant training-derived centroid was subtracted unchanged from both measured and predicted test Δ profiles before recomputing their Pearson correlation.

Centroid accuracy measured whether a prediction was closer to its matched response than to other compounds measured in the same target cell line and dose. For prediction 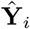 among *n* test observations in its context, we calculated

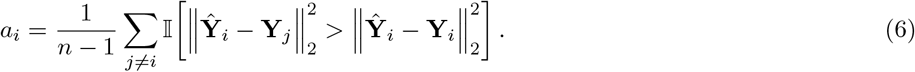

*a*_*i*_ is therefore the fraction of alternative compound responses ranked farther from the prediction than its matched response.

### 4.7 Reproducibility

#### Replicate reproducibility

Profiles were grouped by cell line, drug, dose, and exposure time. We formed unordered pairs within each group, separating profiles measured on the same plate (intra-plate) from those measured on different plates (inter-plate). For Δ-profile comparisons, each treated replicate was first corrected using the mean DMSO profile matched to its own cell line, plate, and exposure time. Figure 3a reports the resulting individual pairwise Pearson correlations. To limit computation, LINCS pairs were randomly subsampled to at most 200 per condition. For analyses requiring one reproducibility estimate per condition, we averaged the Pearson correlations across its available replicate pairs:

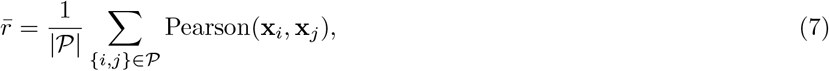

where *p*is the relevant set of replicate pairs. For the dose comparison in Fig. 3a, response magnitude was computed as the arithmetic mean of the Euclidean norms of the unique replicate Δ profiles in each dose–time group.

#### Reproducibility–performance association

Each LINCS test observation was matched by cell line, compound, dose, and exposure time to its mean inter-plate Δ-profile reproducibility. Observations without a defined inter-plate estimate were excluded. For Fig. 3c,d, matched observations were grouped into 0.1-wide reproducibility intervals from ( *−* 0.4, *−* 0.3] through (0.8, 0.9]; bins containing at least 20 observations were retained. We calculated Spearman correlation across all matched, unbinned observations. The reported *R*^2^ was obtained from an unweighted linear regression of the retained bins’ mean prediction correlations on their midpoints.

#### Training data filtering

The reproducibility-filtering experiment reused the fixed LINCS compound split described above. Condition-level inter-plate Δ reproducibility was estimated from all available raw profiles before predictive splitting. Filtering was then optionally applied to training observations. We evaluated thresholds {none, 0.0, 0.1, 0.2, 0.3, 0.4, 0.5*}*, where a numerical threshold retained training observations with a defined mean inter-plate Pearson correlation at or above that value. Each experiment received an independent hyperparameter sweep and was selected by validation Spearman as described above. Test observations were binned by reproducibility, with 0.1 intervals. Bins containing at least 100 observations were retained, yielding 11 intervals from (*−*0.3, *−*0.2] through (0.7, 0.8] in Fig. 3e.

#### Sequencing depth

Sequencing depth was defined as the total UMI count of each Tahoe-100M pseudobulk before library-size normalization and log transformation. The analysis included treated profiles from the main compound-held-out test set. UMI counts were first averaged across profiles contributing to each test observation and then across observations within each target cell line, excluding DMSO profiles. Prediction performance was the mean per-observation Pearson correlation within the same cell line, and its association with mean sequencing depth was assessed using Spearman correlation (Fig. 3f).

### 4.8 Reference-cell analyses

#### Reference-cell choice

We repeated BioPert training with each reference cell line shown in Fig. 4a,b. Each choice defined its own eligible observations by requiring a matched reference response, and the selected reference cell line was excluded from the prediction targets. Models were tuned independently.

#### Reference–target response similarity

For each pair of reference and target cell lines in Tahoe-100M, replicate Δ profiles were first averaged across plates within each compound, dose, and exposure-time condition. We calculated the gene-wise Pearson correlation between the reference and target responses for every shared condition and averaged these correlations with equal weight, excluding pairs with fewer than two shared conditions. This descriptive analysis used all shared conditions in the filtered Tahoe-100M data, including training and held-out compounds. Each point in Fig. 4d represents one reference–target pair, and the association with prediction performance for the target cell line was assessed using Spearman correlation.

### 4.9 Expression variance decomposition

To quantify biological and technical sources of expression variability, we fit separate random-intercept models to each gene using `variancePartition` [24], which estimates variance components by restricted maximum likelihood through `lme4.` The dataset-specific models were

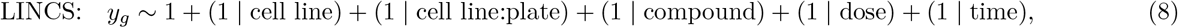

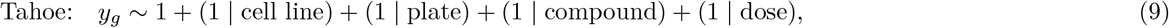

where plate identity was nested within cell line for LINCS and the time term was omitted from Tahoe, as all profiles have the same exposure time. Models were fitted to individual profiles, separately for absolute expression and Δ expression. The absolute-profile analysis included treated and DMSO profiles. For each factor, we report the median across genes of its estimated variance fraction, expressed as a percentage.

### 4.10 Pathway analysis

#### Gene-set enrichment analysis (GSEA)

We performed preranked GSEA [21] for different pairs of cell line and compound on Tahoe-100M. For each case, three profiles were analyzed independently: the measured target cell line response, the corresponding BioPert prediction, and the measured response in the A549 reference cell line. Log-normalized pseudobulk Δ profiles were ranked using all 19,020 protein-coding genes. Exact ties were resolved deterministically using a negligible perturbation. Gene sets were retrieved from the Enrichr public mirrors [51] using `gseapy.get_library(organism=“Human”).` We used the MSigDB_Hallmark_2020 collection (50 gene sets) and `Reactome_2022` collection (1,816 gene sets) [52, 53]. Preranked enrichment was run with GSEApy v1.3.0 [54] using weighted scores (weight 1), gene-set sizes of 15–500 genes, 5,000 gene-set permutations, and seed 42. GSEApy false-discovery-rate (FDR) *q* values were calculated separately for each case, profile, and collection.

For each case and gene-set collection, agreement with the measured target profile was quantified for the reference and prediction using the Spearman correlation between normalized enrichment scores (NES), the mean absolute NES difference across shared pathways, and the proportion of target-significant pathways with matching NES signs. A pathway was classified as target-specific when it had target FDR *q <* 0.05 and either its reference NES had the opposite sign or| NES_reference_ *−* NES_target_ |*≥* 1.0. The **C32-cobimetinib** case was selected for analysis for the maximal divergence between the target C32 and the fixed reference A549, in both oncogenic driver (BRAF-V600E and KRAS-mutant, respectively) and cell lineage (melanocytic and alveolar epithelial, respectively), combined with the well-characterized, directionally predictable mechanism of action of cobimetinib in lines carrying these mutations.

### 4.11 AI tools

OpenAI Codex and Anthropic Claude were used as assistive tools to review and refine analysis code and manuscript text.

## Data availability

The data that support the findings of this study are openly available in the HGNC reference database at https://www.genenames.org/download/statistics-and-files/ (accessed 2025-11-24, see [32]); tahoebio/Tahoe-100M at https://huggingface.co/datasets/tahoebio/Tahoe-100M; LINCS L1000 Phase II at https://clue.io/releases/data-dashboard; the Hallmark and Reactome sets at https://maayanlab.cloud/Enrichr/.

## Code availability

All code used in this study, including scripts for downloading and processing the Tahoe-100M and the LINCS L1000 datasets, is available in the Biopert GitHub repository (https://github.com/lemieux-lab/biopert).

## Acknowledgment

This work was made possible through computational resources provided by Calcul Quebec (calculquebec.ca), the Digital Research Alliance of Canada (alliancecan.ca), and Mila (mila.quebec).

## Author contributions

Conceptualization: L.K., L.L.B., Q.F., S.L. ; Methodology: L.K., L.L.B., Q.F., S.L. ; Software: L.K., L.L.B. ; Validation: L.K., L.L.B. ; Formal analysis: L.K., L.L.B., E.C.B ; Investigation: L.K., L.L.B. ; Data curation: L.K., L.L.B. ; Writing - original draft: L.K., L.L.B., E.C.B ; Writing - review and editing: L.K., L.L.B., Q.F., S.L. ; Visualization: L.K., L.L.B. ; Supervision: Q.F., S.L. ; Project administration: L.K., L.L.B. ; Funding acquisition: Q.F., S.L.

## Funding

This work was supported by the Natural Sciences and Engineering Research Council of Canada [RGPIN-2022-04260 to SL]. This project was undertaken thanks to funding from IVADO and the Canada First Research Excellence Fund. We acknowledge the Fonds de recherche du Québec (FRQ) – secteur Santé for its support of the Institute for Research in Immunology and Cancer (IRIC), an FRQ-designated research center (https://doi.org/10.69777/339366).

## Competing interests

The authors declare no competing interests.

1 https://github.com/theislab/cpa/blob/main/cpa/_utils.py

## Notes

### Competing Interest Statement

The authors have declared no competing interest.

