## Supplemental for "A biological-response compound representation allows chemical perturbation prediction across cell lines"

### Supplementary Information

#### Supplementary Methods

##### Datasets, observations, and analysis cohorts

We used Tahoe-100M and LINCS L1000 as complementary settings for evaluating cross-context chemical perturbation prediction. Tahoe-100M contains single-cell measurements from a fully crossed MOSAIC co-culture design, whereas LINCS contains bulk, high-throughput measurements collected across a larger but more sparsely sampled set of compounds, cell lines, doses, and exposure times. Following the preprocessing described in Methods, the Tahoe-100M data contained 379 compounds and 48 cell lines. Two cell lines, NCI-H2122 and NCI-H596, were removed because more than 90% of their pseudobulks contained fewer than 35,000 UMIs. The processed LINCS data contained 26,426 compounds and 222 cell lines. Supplementary Table 1 reports the final inventory.

One model observation, or condition, represented a combination of target cell line, compound, dose and exposure time. We calculated $\Delta$ profiles for each treated profile by subtracting the unweighted mean of DMSO profiles matched by cell line and plate. We then averaged replicate $\Delta$ profiles across plates within each condition. The target cell input representation was the unweighted mean of all DMSO profiles available for that cell line. This design assumes that an untreated expression profile from the target cell line is available when predicting its treated response.

BioPert additionally requires at least one reference cell $\Delta$ profile matching the compound, dose, and exposure time of the target observation. For a held-out compound, the measured reference cell response was therefore available as an input at inference, although no target cell response for that compound entered model training. Molecular and random representations did not require a matched reference cell measurement, which yielded a larger LINCS training cohort (369,199 observations) than BioPert (225,562 observations). All representation comparisons used the common intersection of eligible test observations.

The default BioPert cohort contained 52,712 Tahoe-100M observations and 294,172 LINCS observations. The train, validation, and test partitions contained 39,685, 5,260, and 7,767 observations for Tahoe-100M and 225,562, 28,630, and 39,980 observations for LINCS, respectively.

| | Tahoe-100M | LINCS |
|---|---|---|
| Untreated profiles (DMSO) | 1,337 | 75,705 |
| Treated profiles | 61,926 | 1,221,534 |
| Profile length (genes) | 19,020 | 978 |
| Cell lines | 48 | 222 |
| Drugs | 379 | 26,426 |
| Doses | 3 | 65 |
| Exposure times | 1 | 11 |
| Plates | 14 | 5,146 |

**Supplementary Table 1:** Dataset inventory after preprocessing.

##### Molecular and random treatment representations

The structure–activity analysis considered 25 compound representations: ten molecular fingerprints, 14 pretrained molecular representations, and one random control. The fingerprints comprised Morgan/ECFP6, RDKit path, and atom-pair fingerprints at 512, 1,024, and 2,048 bits, together with MACCS keys. Morgan fingerprints were binary, non-feature fingerprints with radius 3 and otherwise used RDKit defaults. The pretrained representations comprised CheMeleon, bert-base-smiles, ChemBERTa-5M-MLM, ChemBERTa-5M-MTR, ChemBERTa-77M-MLM, ChemBERTa-100M-MLM, MoLFormer-XL-both-10pct, MolGen-large, Uni-Mol1 (84M), and Uni-Mol2 (84M, 164M, 310M, 570M, and 1.1B). The native representation dimensions ranged from 384 to 2,048.

For Hugging Face transformer models, the extraction code used `pooler_output` when available and otherwise calculated an unweighted mean of the final hidden layer. MolGen input SMILES were converted to SELFIES before embedding. CheMeleon used the fingerprint produced by the released message-passing model.

The random control assigned each exact SMILES a fixed 512-dimensional vector sampled independently from a standard normal distribution. The same vector was reused for a compound throughout training and evaluation. For the structure–activity analysis, fingerprint dissimilarity was measured using Tanimoto distance and continuous-embedding dissimilarity using cosine distance, so larger values consistently represented less similar compounds.

##### Reference-response construction

The default BioPert representation used A549 as the reference cell line. For each target observation, we sampled one of the available reference-cell $\Delta$ profiles matching its compound, dose, and exposure time, and held that sampled profile fixed during training and evaluation. Sampling occurred separately for each emitted target observation. Consequently, target cell lines sharing a treatment did not necessarily receive the same reference replicate. In the resampling ablation, a new matching reference replicate was drawn for each training observation at every epoch, whereas validation and test observations retained their original fixed draws. In the averaging ablation, all matching reference replicates were averaged with equal weight across plates. The three reference-handling strategies used the same test observations and were tuned independently.

For the reference-choice analysis, we repeated model fitting with A549, A-172, COLO 205, and PANC-1 in Tahoe-100M and with A549, MCF7, and PC3 in LINCS. Each reference choice defined its own eligible treatment cohort and was excluded from the prediction targets. We compared references across the target cell lines represented under every reference choice.

##### Structure–activity analysis

We evaluated whether molecular proximity predicted similarity between measured transcriptional responses. Within each condition assay group, we retained one randomly selected $\Delta$ profile for each exact SMILES and formed every unordered pair of compounds. Thus, one analysis row represented a compound pair observed in one shared assay context. Conditions with more compounds contributed more rows because the association was calculated across condition-repeated pairs rather than unique compound pairs. In LINCS, the analysis retained conditions with a mean inter-plate $\Delta$-profile reproducibility of at least 0.7, and conditions without a defined inter-plate estimate were excluded. We calculated Pearson correlation between the molecular distance and the gene-wise Pearson correlation of the two measured $\Delta$ profiles. Confidence intervals resampled compound-pair rows and recalculated the association in each bootstrap sample.

##### Model optimization and selection

All MLPs used ReLU hidden activations and a linear output layer, without dropout, normalization layers, or residual connections. The RMSE objective weighted every gene–observation element equally. PCA used training-set centering without variance scaling and was fitted to the complete training-observation matrix.

We evaluated each checkpoint every ten epochs and retained the epoch with the highest mean validation Spearman correlation within a trial. We then selected the trial whose retained checkpoint obtained the highest validation Spearman correlation. The selected checkpoint was evaluated directly on the test set without retraining on the combined training and validation data. Secondary analyses, including dose encoding, the landmark-gene restriction, reproducibility filtering, cell line holdout, reference choice, and reference handling, received independent hyperparameter searches.

##### Systema sensitivity analysis

For each Tahoe target cell line $l$, we estimated a generic perturbation centroid from training responses, $c_l = |T_l|^{-1} \sum_{i \in T_l} Y_i$. We also estimated dose-specific centroids, $c_{l,d} = |T_{l,d}|^{-1} \sum_{i \in T_{l,d}} Y_i$, from training observations sharing target cell line $l$ and dose $d$. Each training-derived centroid was subtracted unchanged from the corresponding measured and predicted test profiles.

For centroid accuracy, we compared each prediction $\hat{Y}_i$ with every measured test response in the same context using squared Euclidean distance. Accuracy was the fraction of alternative compounds farther from the prediction than its matched response. Equal-distance ties were not counted as outranked alternatives, and contexts containing fewer than two test candidates had undefined accuracy. Confidence intervals resampled test compounds and retained all associated observations.

##### Reproducibility and measurement-quality analyses

Replicate analyses used individual unordered profile pairs. For $\Delta$-profile reproducibility, each treated profile was first corrected using the mean DMSO profile matched to its cell line, plate, and exposure time. Pairs were then classified as intra-plate or inter-plate. LINCS distinct conditions were randomly subsampled to at most 200 pairs. For condition-level analyses, pairwise Pearson correlations were averaged within each condition.

We matched each LINCS test observation to the mean inter-plate reproducibility of its exact condition and excluded observations without an estimate. For descriptive visualization, we used 0.1-wide bins from $(-0.4, -0.3]$ through $(0.8, 0.9]$

and retained bins containing at least 20 observations. The Spearman association used all 29,547 matched, unbinned observations. The reported $R^2$ was obtained from an unweighted linear regression of the 13 retained bin means on their bin midpoints.

The reproducibility-filtering analysis reused the main compound split and changed only the training set. Thresholds of 0.0–0.5 retained conditions whose mean inter-plate reproducibility met or exceeded the threshold. Reproducibility was estimated from the complete raw dataset before predictive splitting, whereas filtering was applied only to training observations. Validation and test observations remained fixed across thresholds, and every threshold received an independent model search. No downsampling control was used, so increasing the threshold jointly changed measurement reproducibility, training-set size, and treatment coverage. Test results were summarized in 0.1-wide reproducibility bins containing at least 100 observations.

For Tahoe-100M, sequencing depth was the total UMI count of each treated pseudobulk before library-size normalization and log transformation. We first averaged depth across profiles contributing to each test observation and then across observations within each target cell line. The association used the mean test Pearson correlation and mean sequencing depth of each of the 47 target cell lines.

##### Cell line generalization and reference–target similarity

We tested generalization to unseen cell lines with independently assigned training, validation, and test cell line sets in addition to the compound split. The observed-compound setting evaluated observations from test cell lines whose compounds occurred in training in other cell lines. The joint setting evaluated the intersection of independently held-out compounds and held-out target cell lines. In both settings, the full-dataset DMSO mean of the held-out target cell line and the matched response from the reference cell line remained available as inputs. We calculated the gain over the copied-reference baseline for each observation, averaged gains within target cell lines, and resampled target cell lines for confidence intervals.

For each Tahoe reference–target pair, we averaged replicate $\Delta$ profiles by cell line, compound, dose, and exposure time, calculated gene-wise Pearson correlation for every shared condition, and then averaged conditions with equal weight. Pairs required at least two shared conditions. This descriptive analysis used all shared conditions in the filtered Tahoe-100M data, including training and held-out compounds. The association between mean response similarity and prediction performance treated each of the 188 pairs of reference and target cell lines as one observation.

##### Gene-set enrichment analysis

We performed preranked GSEA for ten Tahoe-100M test cases selected from the 7,767 candidate observations. For each case, we analyzed the measured target-cell response, the BioPert prediction, and the measured A549 reference response separately. All 19,020 protein-coding genes were ranked by their signed $\Delta$-profile values after mapping Tahoe gene tokens to uppercase HGNC symbols. We resolved exact ties deterministically using a negligible perturbation. We used the MSigDB Hallmark 2020 and Reactome 2022 collections obtained through the Enrichr public mirrors. GSEApy v1.3.0 was run with weighted scores, gene-set sizes of 15–500 genes, 5,000 gene-set permutations, and seed 42. The size filter retained all 50 Hallmark sets and 1,177 Reactome sets. FDR $q$ values were calculated separately for each case, profile, and collection. Pathway-level concordance was summarized using Spearman correlation and mean absolute error between normalized enrichment scores. Direction agreement was evaluated among pathways significant in the measured target at FDR $q \leq 0.05$, and leading-edge overlap was summarized using the Jaccard index.

##### Statistical analysis

Unless stated otherwise, confidence intervals used 1,000 bootstrap resamples and the 2.5th and 97.5th percentiles. Main pairwise model comparisons used two-sided paired Wilcoxon signed-rank tests on matched per-observation correlations. Benjamini–Hochberg correction was applied separately within Tahoe-100M and LINCS to the 18 model-versus-random comparisons and across the 11 reproducibility-bin comparisons of unfiltered training with the 0.5 threshold.

##### Hyperparameters

- batch size: $\{32, 64, 128, 256\}$
- target learning rate: log-uniform over $[10^{-6}, 10^{-2}]$
- weight decay: $\{0, 10^{-4}, 10^{-3}, 10^{-2}, 10^{-1}\}$

- hidden-layer dimensions: `[512,512,512]`, `[1024,1024]`, and `[1024,512,256]`; `[1024,128,1024]`, `[512,256,512]`, `[256,512,256]`, and `[128,1024,128]`
- maximum number of training epochs: $\{100, 200, 500\}$.
- number of PCA components for the untreated target cell line: $\{32, 64, 128, 256, 1024\}$ (1024 omitted from LINCS sweeps)
- number of PCA components for the treatment representation: $\{0, 32, 64, 128, 256\}$ (0 retained the native molecular representation)

| Configuration | LR | BSZ | Weight decay | Hidden layers | Max epochs | $n_{\text{PCA}}^{\text{expr}}$ | $n_{\text{PCA}}^{\text{molec}}$ | Epoch |
|---|---|---|---|---|---|---|---|---|
| ECFP6-512 | 2.29e-04 | 32 | 0.1 | [1024, 1024] | 500 | 32 | — | 180 |
| ECFP6-1024 | 1.44e-05 | 128 | 0.1 | [1024, 1024] | 200 | 128 | 64 | 180 |
| ECFP6-2048 | 5.68e-05 | 32 | 0.1 | [1024, 1024] | 500 | 64 | 64 | 30 |
| BioPert | 7.08e-05 | 32 | 0.1 | [1024, 1024] | 500 | 128 | 128 | 30 |
| CheMeleon | 1.29e-04 | 256 | 0.0001 | [1024, 1024] | 100 | 256 | — | 20 |
| ChemBERTa-100M-MLM | 2.61e-05 | 32 | 0.1 | [512, 512, 512] | 500 | 64 | — | 30 |
| ChemBERTa-5M-MLM | 1.37e-05 | 256 | 0.1 | [1024, 1024] | 200 | 128 | 64 | 180 |
| ChemBERTa-5M-MTR | 2.01e-05 | 32 | 0.1 | [1024, 1024] | 200 | 64 | 256 | 90 |
| ChemBERTa-77M-MLM | 2.75e-05 | 64 | 0 | [1024, 1024] | 200 | 256 | 256 | 50 |
| MoLFormer-XL | 1.38e-04 | 256 | 0.1 | [1024, 1024] | 200 | 256 | 256 | 20 |
| MolGen-large | 3.25e-05 | 128 | 0 | [1024, 1024] | 200 | 128 | 64 | 40 |
| Random | 3.58e-05 | 32 | 0.1 | [1024, 1024] | 200 | 256 | 32 | 20 |
| UniMolv1-84M | 1.80e-05 | 256 | 0.0001 | [1024, 1024] | 500 | 64 | 32 | 60 |
| UniMolv2-1.1B | 3.37e-04 | 64 | 0.1 | [1024, 128, 1024] | 500 | 64 | — | 160 |
| UniMolv2-164M | 2.23e-04 | 32 | 0.001 | [512, 256, 512] | 100 | 128 | — | 40 |
| UniMolv2-310M | 2.94e-04 | 32 | 0.1 | [1024, 128, 1024] | 100 | 256 | — | 80 |
| UniMolv2-570M | 2.72e-05 | 128 | 0.1 | [1024, 128, 1024] | 500 | 256 | — | 100 |
| UniMolv2-84M | 3.58e-05 | 64 | 0 | [1024, 128, 1024] | 200 | 256 | — | 80 |
| bert-base-smiles | 6.53e-05 | 128 | 0.01 | [1024, 1024] | 500 | 256 | 32 | 20 |

**Supplementary Table 2:** Best-by-validation-Spearman hyperparameters, LINCS.

| Configuration | LR | BSZ | Weight decay | Hidden layers | Max epochs | $n_{\text{PCA}}^{\text{expr}}$ | $n_{\text{PCA}}^{\text{molec}}$ | Epoch |
|---|---|---|---|---|---|---|---|---|
| ECFP6-512 | 5.72e-05 | 256 | 0.1 | [512, 256, 512] | 200 | 256 | 32 | 170 |
| ECFP6-1024 | 2.14e-05 | 128 | 0.0001 | [512, 256, 512] | 500 | 256 | 32 | 220 |
| ECFP6-2048 | 7.91e-05 | 256 | 0.1 | [1024, 128, 1024] | 200 | 64 | 32 | 120 |
| BioPert | 8.91e-04 | 256 | 0.001 | [1024, 128, 1024] | 200 | 128 | 128 | 190 |
| BioPert (landmark) | 2.33e-04 | 256 | 0.1 | [1024, 1024] | 500 | 64 | 64 | 220 |
| CheMeleon | 4.11e-06 | 128 | 0 | [512, 512, 512] | 500 | 32 | 32 | 400 |
| ChemBERTa-100M-MLM | 1.13e-05 | 64 | 0.001 | [256, 512, 256] | 500 | 32 | 32 | 130 |
| ChemBERTa-5M-MLM | 6.95e-06 | 32 | 0.001 | [256, 512, 256] | 500 | 256 | 32 | 190 |
| ChemBERTa-5M-MTR | 1.18e-05 | 64 | 0.01 | [1024, 1024] | 200 | 1024 | 32 | 150 |
| ChemBERTa-77M-MLM | 6.57e-06 | 32 | 0.1 | [1024, 128, 1024] | 500 | 1024 | 32 | 480 |
| MoLFormer-XL | 1.10e-05 | 32 | 0.0001 | [1024, 1024] | 100 | 128 | 32 | 50 |
| MolGen-large | 1.69e-05 | 128 | 0 | [1024, 1024] | 200 | 128 | 32 | 140 |
| Random | 2.19e-05 | 256 | 0.0001 | [512, 512, 512] | 200 | 128 | 32 | 150 |
| UniMolv1-84M | 1.62e-05 | 64 | 0.1 | [1024, 1024] | 200 | 256 | 32 | 90 |
| UniMolv2-1.1B | 5.36e-06 | 64 | 0.01 | [512, 512, 512] | 100 | 128 | — | 100 |
| UniMolv2-164M | 9.02e-04 | 128 | 0.1 | [1024, 1024] | 500 | 64 | — | 30 |
| UniMolv2-310M | 1.47e-04 | 32 | 0.1 | [1024, 512, 256] | 200 | 128 | — | 10 |
| UniMolv2-570M | 5.34e-03 | 256 | 0.1 | [512, 512, 512] | 100 | 32 | — | 40 |
| UniMolv2-84M | 2.84e-05 | 32 | 0.01 | [1024, 1024] | 200 | 256 | — | 20 |
| bert-base-smiles | 3.22e-05 | 64 | 0.01 | [256, 512, 256] | 100 | 64 | 32 | 70 |

**Supplementary Table 3:** Best-by-validation-Spearman hyperparameters, Tahoe.

### Supplementary Data

#### Dataset characteristics and measurement variation

The two datasets differed in both assay scale and the distribution of treatment-induced expression changes. Across all genes and treated profiles, $\Delta$ values were centered near zero (Tahoe-100M: mean 0.0006, s.d. 0.073; LINCS: mean $-0.0065$, s.d. 0.700). LINCS had a broader range ($-14.6$ to 14.4) than Tahoe-100M ($-3.8$ to 5.2), consistent with the different measurement technologies and processing.

Variance decomposition further separated biological and technical sources of variation (Supplementary Fig. 1). In absolute profiles, cell line identity accounted for the largest median fraction of gene-level variance in both Tahoe-100M (75.2%) and LINCS (53.3%). Plate identity accounted for 2.4% in Tahoe-100M and 15.7% in LINCS. After DMSO subtraction, the median cell line component decreased to 3.6% in Tahoe-100M and 0.8% in LINCS, whereas residual variation increased to 71.4% and 74.2%, respectively. Drug identity accounted for 16.3% of median gene-level $\Delta$ variance in Tahoe-100M but 5.5% in LINCS. These distributions show that baseline subtraction removes much of the baseline cell line identity while leaving a larger batch effect component in LINCS.

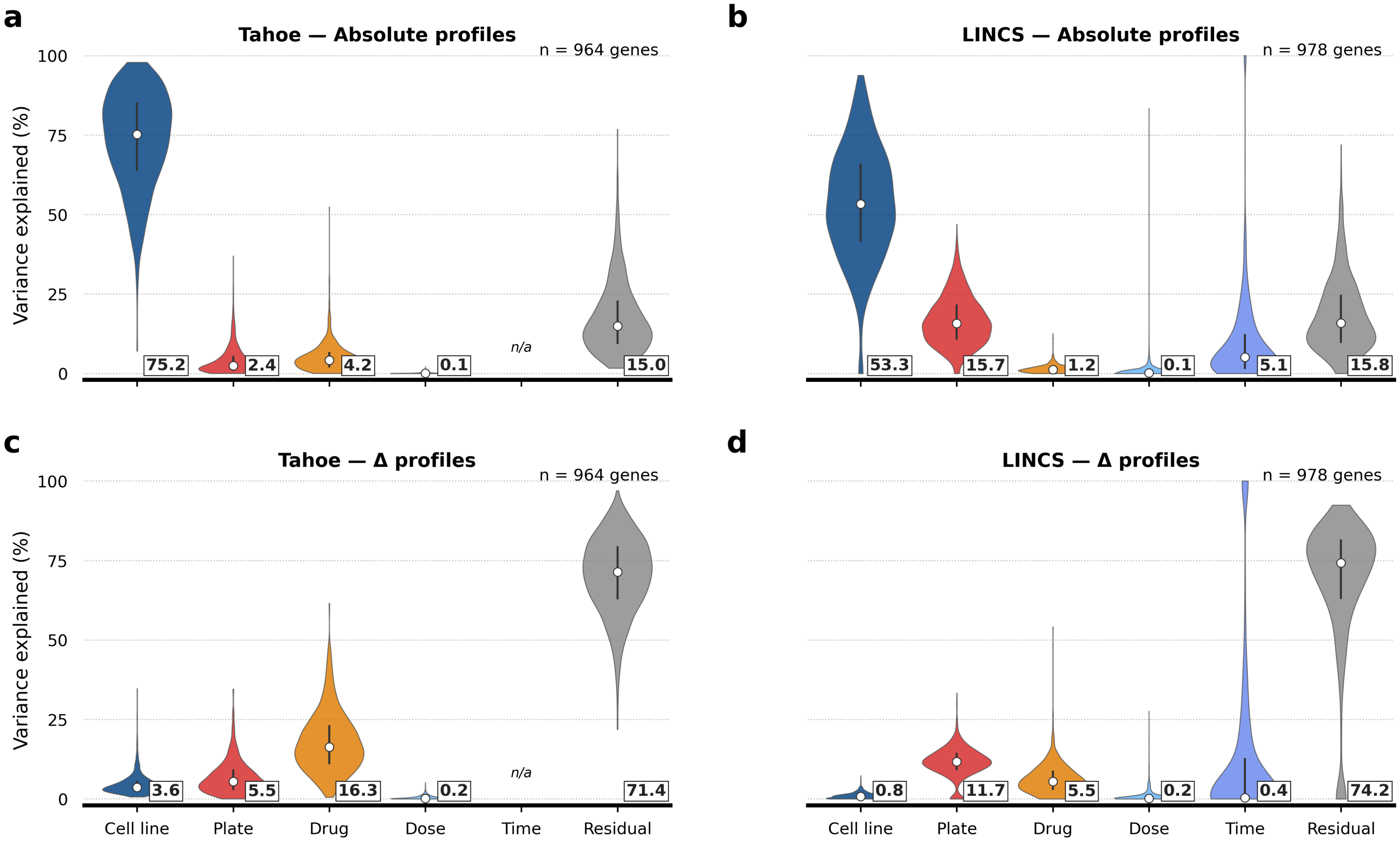


**Supplementary Fig. 1: Expression variance decomposition across experimental and biological factors.** Per-gene variance fractions were estimated using REML mixed-effects models for absolute profiles in Tahoe-100M (**a**) and LINCS (**b**) and for $\Delta$ profiles in Tahoe-100M (**c**) and LINCS (**d**). Violins show the distributions across analyzed genes; white points and adjacent labels indicate medians, and vertical bars indicate interquartile ranges. Tahoe-100M exposure time was constant and was therefore omitted. The analysis contained 964 Tahoe-100M genes and 978 LINCS genes.

Absolute profiles remained highly reproducible across both datasets (Supplementary Fig. 2). Median correlations ranged from 0.982 to 0.994 across the four Tahoe comparisons and from 0.904 to 0.940 across the LINCS comparisons. The lower inter-plate reproducibility in LINCS was present for both untreated and treated profiles, but the reduction was small relative to the deterioration observed after DMSO subtraction.

Within-plate LINCS $\Delta$ reproducibility was strongly associated with dose and exposure time (Supplementary Fig. 3). The 20 $\mu$M, 24-h condition formed a distinct high-reproducibility mode, with a median pairwise Pearson correlation of 0.748 across 35,476 pairs. The 10 $\mu$M, 24-h condition had a median of 0.595 across 20,765 pairs, whereas all remaining conditions had a median of 0.239 across 58,729 pairs.

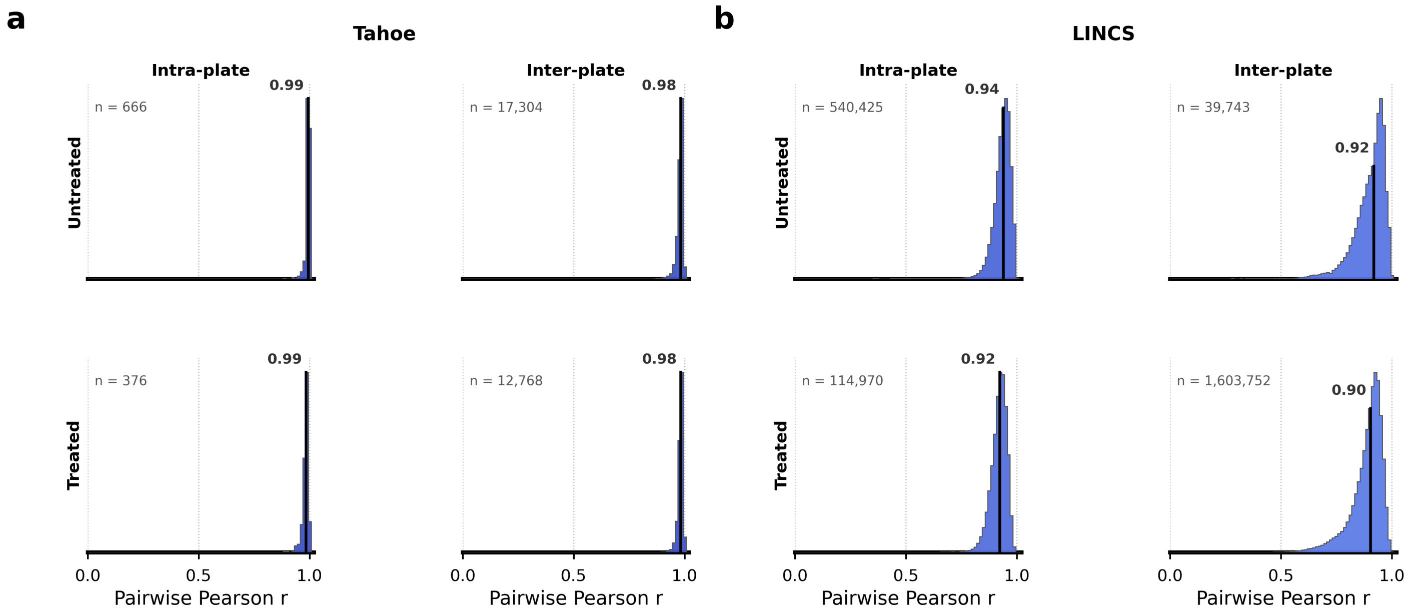


**Supplementary Fig. 2: Absolute expression profiles are reproducible within and across plates.** Distributions show pairwise Pearson correlations for untreated and treated absolute profiles in Tahoe-100M (**a**) and LINCS (**b**), separated into intra-plate and inter-plate comparisons. Vertical lines and labels mark medians; $n$ denotes individual unordered replicate pairs. Tahoe-100M medians were 0.994 (untreated intra-plate, $n = 666$), 0.982 (untreated inter-plate, $n =$ 17,304), 0.985 (treated intra-plate, $n = 376$), and 0.983 (treated inter-plate, $n =$ 12,768). LINCS medians were 0.940 (untreated intra-plate, $n =$ 540,425), 0.918 (untreated inter-plate, $n =$ 39,743), 0.923 (treated intra-plate, $n =$ 114,970), and 0.904 (treated inter-plate, $n =$ 1,603,752).

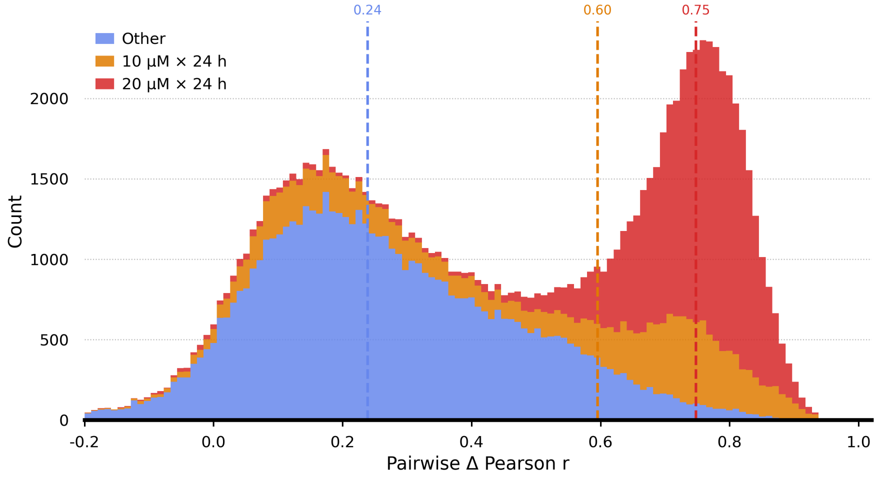


**Supplementary Fig. 3: Dose and exposure time explain the bimodal distribution of intra-plate LINCS Δ-profile reproducibility.** Stacked histograms show individual pairwise Pearson correlations for the 20 $\mu$M, 24-h condition (red), the 10 $\mu$M, 24-h condition (orange), and all other dose–time conditions (blue). Dashed lines indicate the group medians: 0.748, 0.595, and 0.239, respectively.

##### Molecular similarity and predictive robustness

The association between molecular distance and transcriptional-response similarity was weak and representation-dependent (Supplementary Fig. 4). In Tahoe-100M, coefficients ranged from −0.054 for the random control to 0.189 for Uni-Mol v1. In LINCS, molecular distances were negatively associated with response similarity for all displayed representations, but the coefficients remained modest, ranging from −0.017 for the random control to −0.276 for ECFP6.

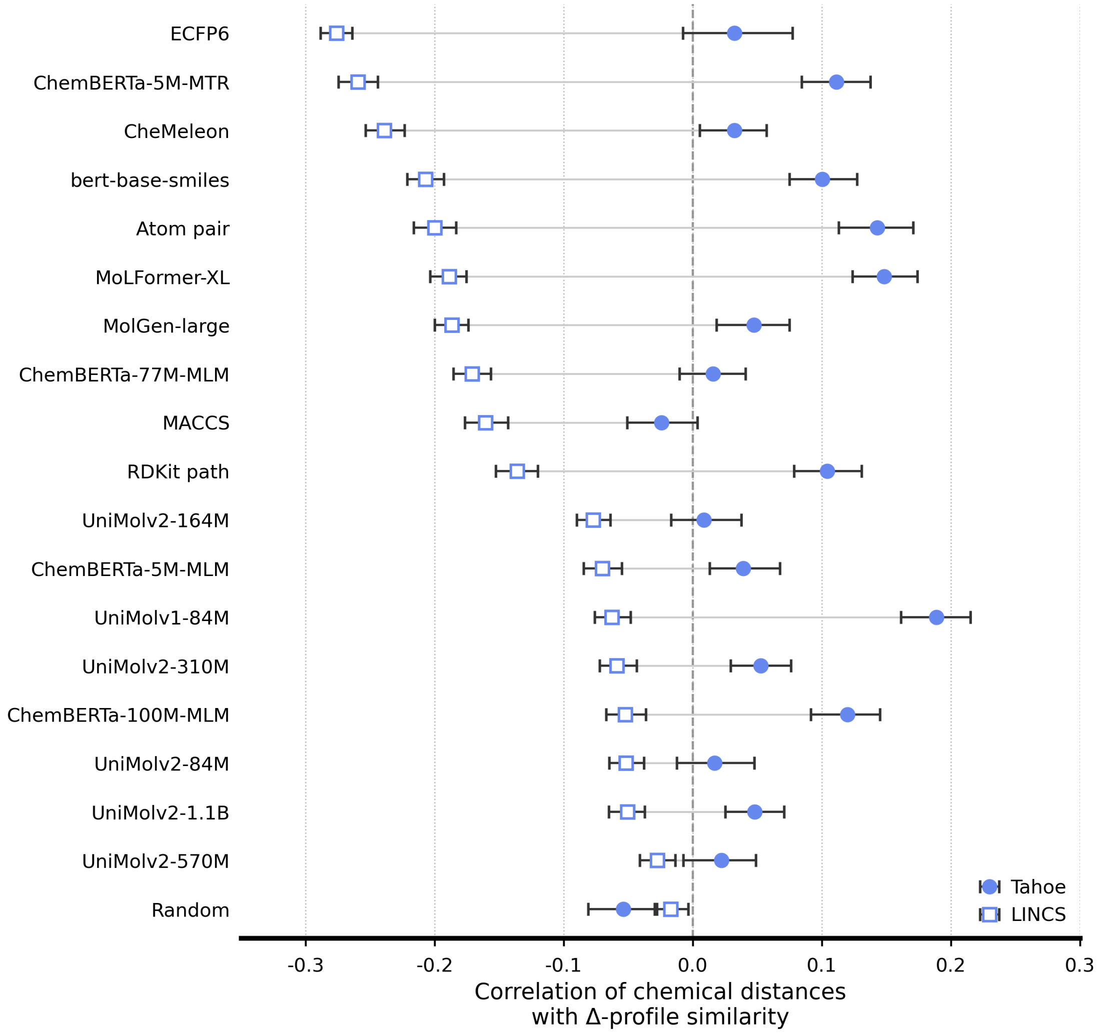


**Supplementary Fig. 4: Association between molecular distance and transcriptional-response similarity across representation families.** Points show Pearson correlations between molecular distance and measured Δ-profile similarity in Tahoe-100M (filled circles; 5,102 compound-pair rows) and LINCS (open squares; 20,016 compound-pair rows). Error bars indicate 95% percentile-bootstrap confidence intervals obtained by resampling pair rows. Fingerprints use Tanimoto distance, and continuous representations use cosine distance.

The principal ranking of predictive methods was robust to the evaluation metric (Supplementary Fig. 5). BioPert obtained the highest Spearman correlation in both datasets and the lowest $\ell_2$ error in Tahoe-100M. In LINCS, $\ell_2$ errors differed little among methods despite clearer separation by correlation, consistent with a setting in which much of the output magnitude is difficult to predict but treatment-specific rank structure remains detectable. Molecular representations remained close to the random control across both correlation- and distance-based metrics.

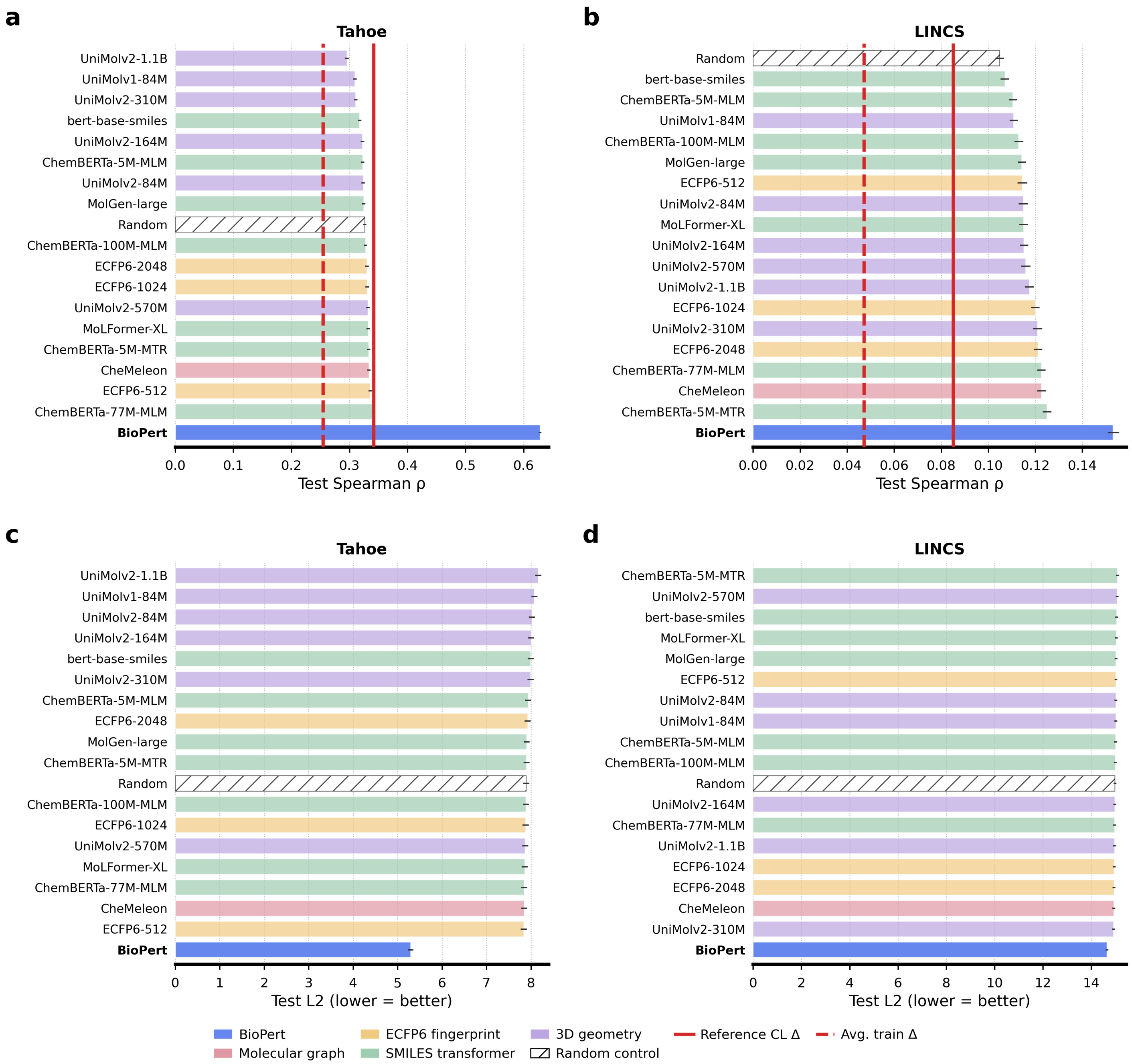


**Supplementary Fig. 5: Representation comparisons are robust to rank-correlation and distance-based evaluation.** Mean test Spearman correlation for Tahoe-100M (**a**) and LINCS (**b**), and mean test $\ell_2$ error for Tahoe-100M (**c**) and LINCS (**d**). Lower $\ell_2$ values indicate better predictions. Bars show validation-selected models evaluated on the common test cohort. Solid and dashed red lines indicate the copied-reference and mean-training-Δ baselines, respectively.

Explicit dose encoding improved validation performance for several molecular representations, particularly in LINCS (Supplementary Fig. 6). The multiplicative gate was the strongest encoding for the random control and each displayed molecular family in LINCS, and was therefore used for subsequent molecular and random-representation comparisons. BioPert did not receive a separate dose or exposure-time encoding because its matched reference response already incorporated these conditions.

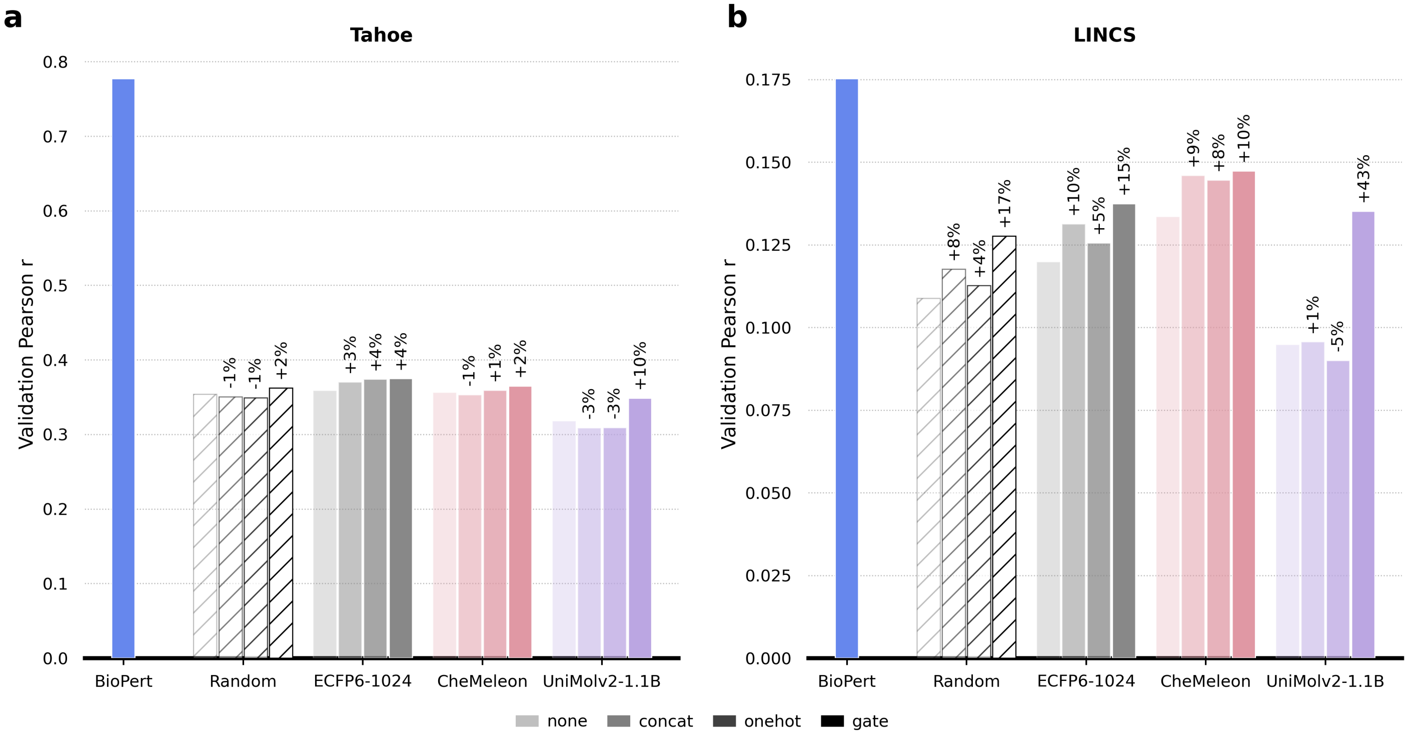


**Supplementary Fig. 6: Validation performance across dose-encoding strategies.** Mean validation Pearson correlation for BioPert and four representative non-biological treatment representations in Tahoe-100M (**a**) and LINCS (**b**). Non-biological representations were evaluated without explicit dose encoding and with concatenation, one-hot encoding, or a learned multiplicative gate. Percentages show the relative difference from the unencoded configuration. BioPert is shown once because the measured reference response already matches dose and exposure time.

Differences among molecular representations were not explained by a monotonic relationship with MLP parameter count, native treatment-representation size, or retained target-cell representation size (Supplementary Fig. 7). BioPert remained separated from the molecular and random controls across a wide range of selected model sizes. This pattern indicates that its performance advantage reflects the information contained in the biological-response input rather than a systematically larger predictor.

Restricting the Tahoe-100M output to the intersection with the LINCS landmark-gene space produced a mean test Pearson correlation of 0.792 for 964 genes, compared with 0.791 in the complete Tahoe-100M output space. The close agreement indicates that BioPert's aggregate performance was not driven by the larger Tahoe-100M gene space.

BioPert also retained substantial performance after removal of generic perturbation centroids. The mean Tahoe-100M test Pearson correlation was 0.791 before centering, 0.733 after subtracting cell-line-specific centroids, and 0.726 after subtracting centroids specific to each cell line and dose. Under the correction defined by cell line and dose, the matched target response was closer to the BioPert prediction than 96.1% of alternative test-compound responses in the same context, compared with the 50% value of the generic centroid baseline. These analyses support compound-specific prediction beyond shared treatment responses.

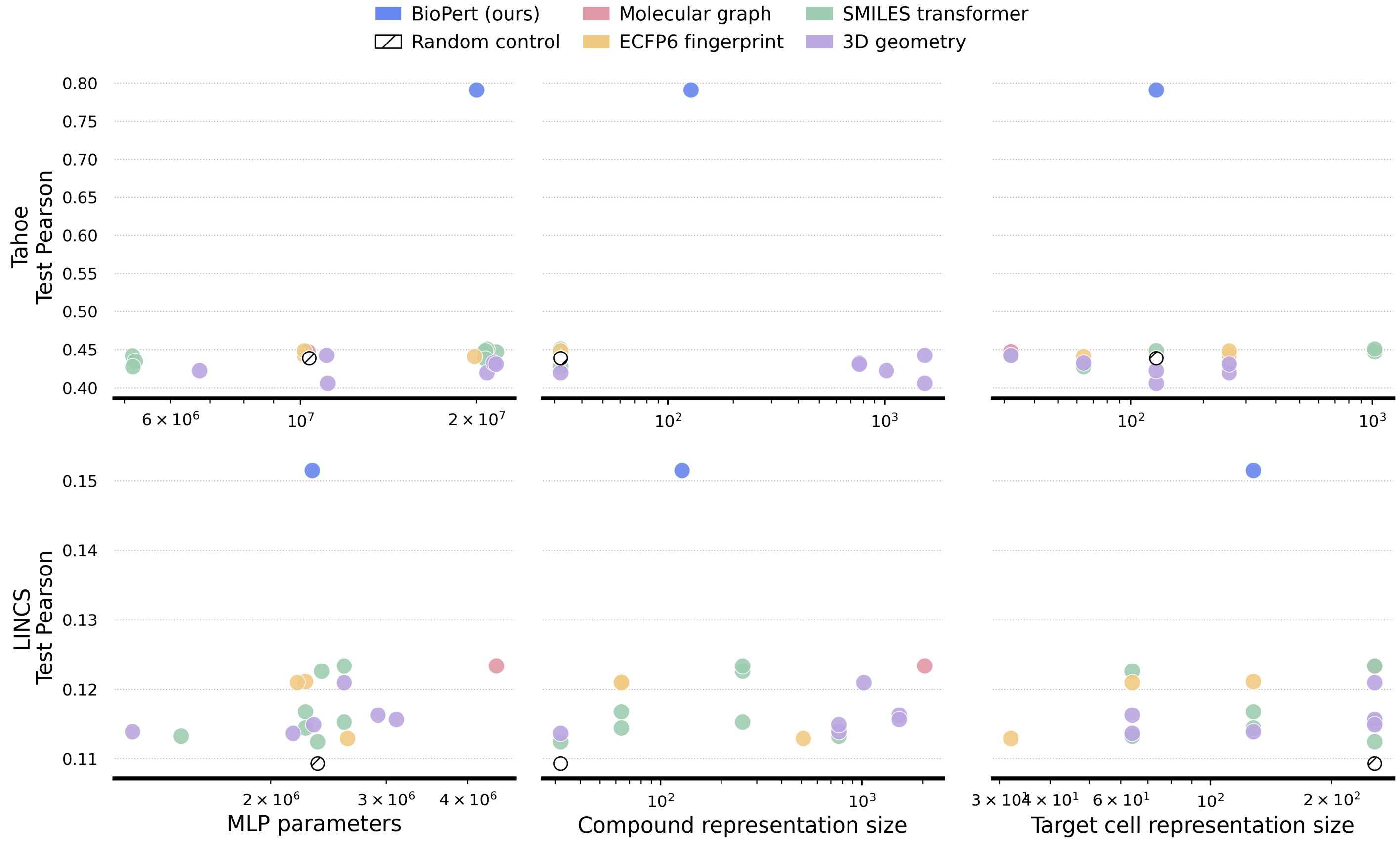


**Supplementary Fig. 7: Predictive performance is not determined by model or representation size.** Test Pearson correlation for the validation-selected Tahoe-100M models (top row) and LINCS models (bottom row) as a function of total MLP parameter count (left), effective treatment-representation dimension after optional PCA (middle), and effective target-cell representation dimension after PCA (right). Each point represents one selected model configuration. Axes for size variables are logarithmic.

#### Reproducibility constrains performance across representations

Prediction performance increased with test-condition reproducibility for every evaluated representation (Supplementary Fig. 8). The curves followed a common sigmoidal pattern: correlations were close to zero for poorly reproducible conditions and rose sharply above inter-plate reproducibility values of approximately 0.2–0.3. BioPert remained the strongest model across most of the reproducibility range, but the convergence of methods in low-reproducibility bins indicates that target noise imposed a shared performance floor.

29,547 test observations had a matched inter-plate reproducibility estimate. Prediction and reproducibility were positively associated across these observations (Spearman $\rho = 0.392$, 95% CI 0.381–0.404). Mean prediction correlation increased from approximately 0.06–0.10 in bins centered near zero to 0.70 in the $(0.7, 0.8]$ bin and 0.71 in the $(0.8, 0.9]$ bin. An unweighted regression through the 13 retained bin means yielded $R^2 = 0.924$.

Filtering the training data did not improve average performance. Raising the minimum training-condition reproducibility to 0.5 reduced the training cohort from 225,562 to 7,492 observations and decreased overall mean test Pearson correlation from 0.151 to 0.084. The filtered model produced small gains in the most reproducible test bins but degraded predictions across the more common low- and intermediate-reproducibility bins. Because the experiment did not include a size-matched downsampling control, it does not separate the effect of reproducibility from the simultaneous loss of training observations and treatment coverage.

In Tahoe-100M, mean sequencing depth varied across the 47 target cell lines and was strongly associated with mean test performance (Spearman $\rho = 0.948$, 95% CI 0.882–0.978). This cell-line-level result complements the LINCS replicate analysis and supports the conclusion that measurement quality constrains achievable prediction accuracy across both assay technologies.

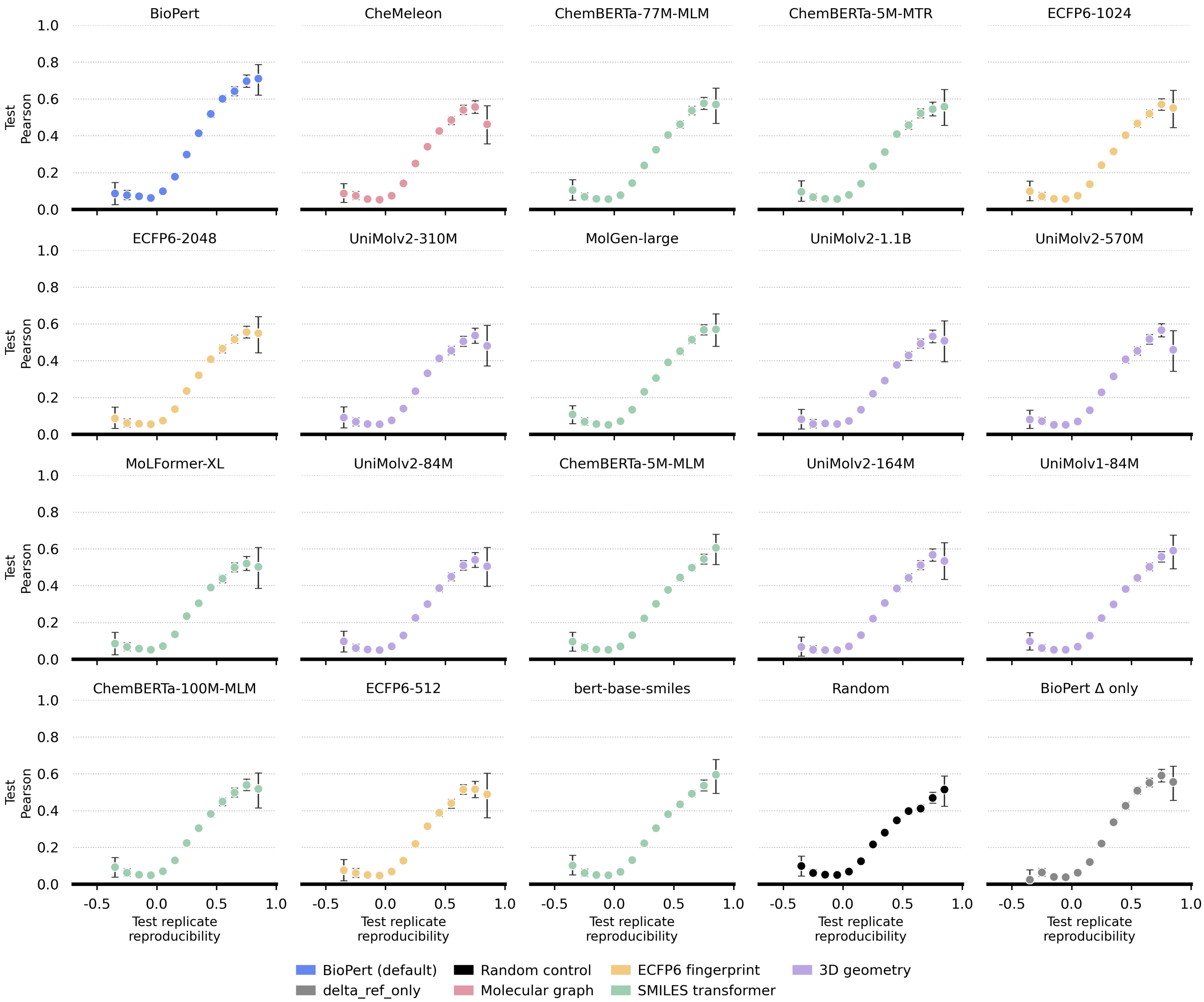


**Supplementary Fig. 8: Prediction accuracy increases with test-condition reproducibility across representation methods.** Mean LINCS test Pearson correlation is plotted against binned inter-plate Δ-profile reproducibility for BioPert, molecular representations, the random control, and the copied-reference baseline. Bins have width 0.1 and contain at least 20 matched test observations.

#### Reference-cell and cellular-context analyses

Changing the reference cell line altered aggregate performance by less than 0.03 within each dataset, but the rank differences among references were detectable when target cell lines were treated as blocks. The Friedman statistics were 111.1 across 44 common Tahoe target cell lines and 61.6 across 179 common LINCS target cell lines. The median spread in target-cell performance across references was 0.028 in Tahoe-100M and 0.020 in LINCS, indicating that reference choice had a modest aggregate effect despite systematic target-specific differences.

Across the 188 Tahoe reference–target pairs, mean response similarity was positively associated with target-cell prediction performance (Spearman $\rho = 0.793$, 95% CI 0.721–0.847). This relationship used both training and held-out compounds and is therefore descriptive rather than an independent test-set-only analysis. Nevertheless, models remained predictive for reference–target pairs with low measured response similarity, supporting transfer beyond near-identical cellular responses.

Reference-profile aggregation improved LINCS predictions. Mean test Pearson correlation was 0.151 for the fixed single-profile strategy, 0.156 with training-time resampling, and 0.178 after averaging all available reference replicates. On the shared 39,980-observation test cohort, averaging improved the mean per-observation correlation by 0.0264 over the fixed strategy (95% CI 0.0246–0.0282), whereas resampling improved it by 0.0042 (95% CI 0.0028–0.0057). These comparisons show that reducing measurement noise in the reference representation was more effective than exposing the model to replicate variability during training.

Holding out target cell lines produced a more difficult generalization setting than holding out compounds alone. Mean gains over the copied-reference baseline decreased to 0.078 in Tahoe-100M and 0.054 in LINCS when compounds were observed during training, but target cell lines were held out. When both compounds and target cell lines were held out independently, the corresponding gains were 0.042 in each dataset. Because these analyses supplied the held-out cell line's full-dataset untreated profile, they test prediction in a new treated context given an observed baseline state rather than fully zero-shot prediction for an unseen cell line.
